# A software tool for at-home measurement of sensorimotor adaptation

**DOI:** 10.1101/2023.12.12.571359

**Authors:** Jihoon Jang, Reza Shadmehr, Scott T. Albert

## Abstract

Sensorimotor adaptation is traditionally studied in well-controlled laboratory settings with specialized equipment. However, recent public health concerns such as the COVID-19 pandemic, as well as a desire to recruit a more diverse study population, have led the motor control community to consider at-home study designs. At-home motor control experiments are still rare because of the requirement to write software that can be easily used by anyone on any platform. To this end, we developed software that runs locally on a personal computer. The software provides audiovisual instructions and measures the ability of the subject to control the cursor in the context of visuomotor perturbations. We tested the software on a group of at-home participants and asked whether the adaptation principles inferred from in-lab measurements were reproducible in the at-home setting. For example, we manipulated the perturbations to test whether there were changes in adaptation rates (savings and interference), whether adaptation was associated with multiple timescales of memory (spontaneous recovery), and whether we could selectively suppress subconscious learning (delayed feedback, perturbation variability) or explicit strategies (limited reaction time). We found remarkable similarity between in-lab and at-home behaviors across these experimental conditions. Thus, we developed a software tool that can be used by research teams with little or no programming experience to study mechanisms of adaptation in an at-home setting.

**Significance:** Sensorimotor learning is traditionally studied in the laboratory, but recent public health emergencies have caused the community to consider at-home data collection. To accelerate this effort, we implemented a software tool that remotely tracks motor learning. Compared with previous remote data collection strategies, our software (1) generates experiments of arbitrary length that (2) run locally on a participant’s laptop which (3) can be modified without any programming expertise in the research laboratory. Here we show a close correspondence between behaviors captured by our tool and those observed in laboratory environments including savings, interference, spontaneous recovery, and variations in implicit and explicit learning due to changes in perturbation variance, reaction time constraints, and feedback delay. Our software and its corresponding manuals are available here: https://osf.io/e8b63/.

## Introduction

The errors we experience during goal-directed movements induce learning, allowing the nervous system to predictively counteract external perturbations or internal changes to our body^1,2^. Sensorimotor learning is overwhelmingly studied within the laboratory: controlled in-lab settings replete with specialized equipment and personnel to monitor experimental progress and participant compliance. For many paradigms, e.g., force-field adaptation^3–7^, an in-lab approach is required due to complex robotic systems needed to deliver physical perturbations, or neuromodulation^8,9^. Controlled laboratory studies are optimal in other ways: e.g., maximizing participant engagement, ensuring clarity of task instructions, and eliminating confounding variables in the experiment environment. But these experiments also pose major limitations: recruitment within academic campuses limits sociodemographic diversity in participant samples, study centers may lack access to patient populations when the university is not associated with a clinical center, and public health concerns such as the recent COVID-19 pandemic can shutter the data collection pipeline.

To address these problems, recent studies^10–12^ have begun to examine the viability of at-home testing using an experimental paradigm that is central to sensorimotor adaptation: visuomotor rotations. In visuomotor rotation studies, the participant controls a cursor’s motion in a 2-dimensional plane either with a robotic manipulandum^13^, a joystick^14^, a tracker on their hand^15,16^, or a stylus on a tablet^17^. At some point, the cursor’s motion is rotated relative to the hand’s actual path, resulting in an error that is countered from one trial to another by parallel subconscious (implicit) and intentional (explicit) adaptive systems^18–20^ (Fig. 1A). Because this paradigm involves visual manipulation of a virtual object, this task is particularly amenable to at-home examination. Indeed, recent laptop-based at-home studies suggest that a few laboratory-based observations such as the response to invariant errors^10^, variations in rotation magnitude^10,12^, and total target count^11^, reproduce within at-home participant samples.

**Figure 1.**
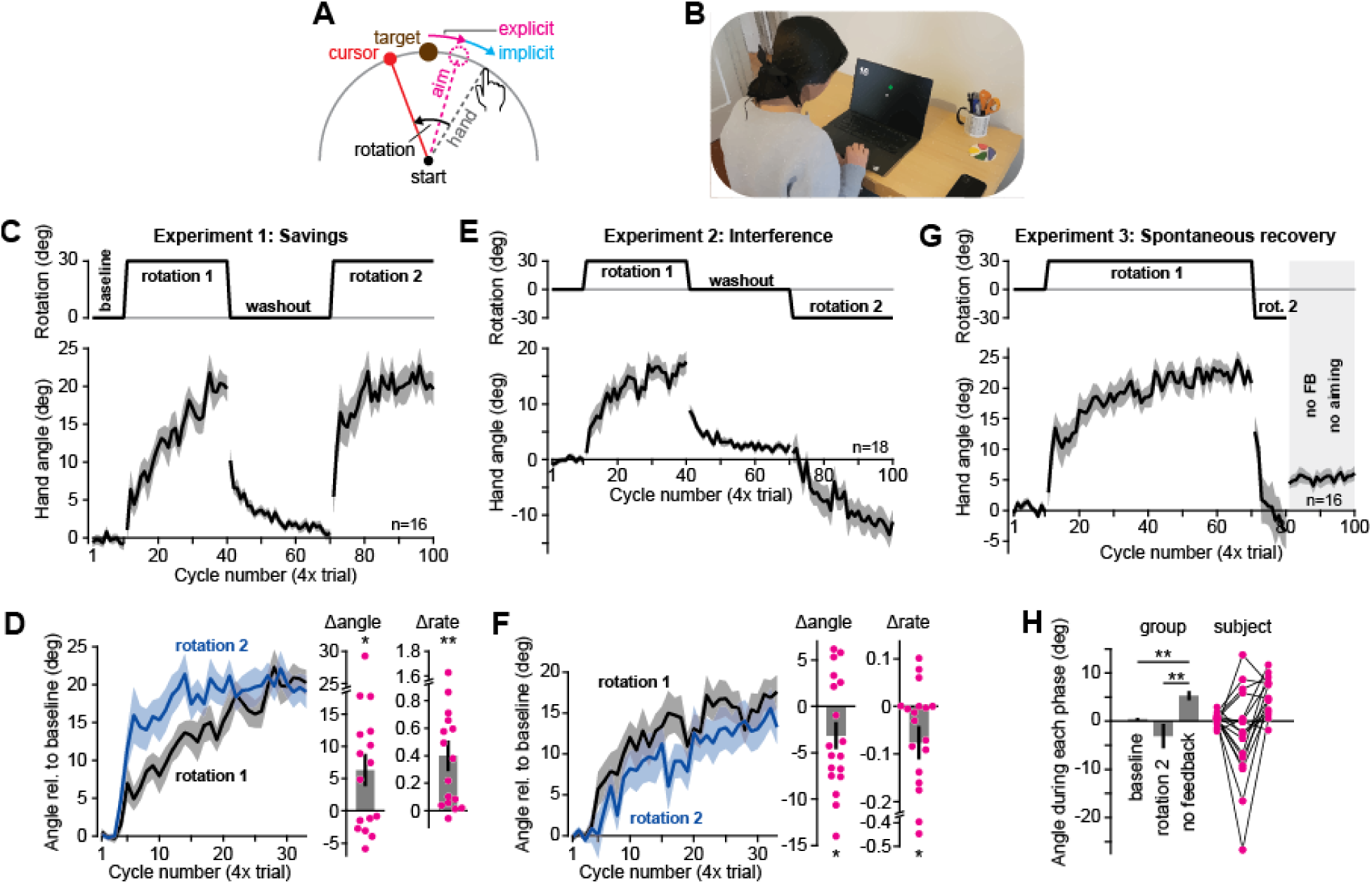
Experience modulates motor adaptation in an unmonitored at-home setting. **A.** Participants controlled a cursor with their hand. A rotation was imposed at times, altering the path of the cursor relative to the hand. Participants adapted, counter-rotating their movement angle. This adaptation is known to consist of explicit (reaiming) and implicit (subconscious adjustment) components. **B.** At-home setup. Participants completed this task at home using their laptop’s trackpad or mouse. **C.** Savings experiment (n=16). The perturbation schedule and movement angles during each block. **D.** Learning curve comparison. Movement angles during each rotation period in **C** were baseline-subtracted (the average angle across the 3 epochs preceding rotation onset were subtracted) and overlaid for visual comparison. To quantify savings, we compared the early movement angle (initial 3 rotation epochs, baseline subtracted) across each rotation period (Δangle, second rotation minus first rotation). We fit a 3-parameter exponential (constrained to ‘begin’ at the pre-rotation angle) and reported the change in growth rate across each rotation exposure (Δrate, second rotation minus first rotation). **E-F.** Same as **C**-**D** but for the anterograde interference paradigm (Experiment 2, n=18). **G.** Spontaneous recovery protocol. This study (n=16) started with a baseline period, rotation 1, brief rotation 2, and no aiming period (no feedback). Movement angles throughout the task are shown at bottom. **H.** To detect spontaneous recovery, average movement angles were calculated over three periods: (1) baseline (cycles 1-10), (2) end of the counter-rotation rotation (cycles 79 and 80), and the no-aiming period (cycles 81-100). Error-bars indicate mean ± SEM. Points in the bar graphs show individual participants. Statistics in **D** and **F** denote a paired t-test. Statistics in **H** denote a repeated-measures ANOVA: *p<0.05, **p<0.01.

Here, we designed a new software tool for these emerging studies, and then tested the tool by asking whether experimental conditions known to increase or reduce adaptation, and its component implicit and explicit systems, could be replicated in the at-home setting. The software runs locally on the participant’s device, thus minimizing potential issues with internet connectivity and hosting services, with audiovisual instructions integrated into the testing environment. We tested the software by focusing on three adaptive phenomena: (1) *savings*, acceleration in learning rate when past perturbations are re-experienced^15,21–24^, (2) *interference*, slowing in adaptation rate when 2 consecutive perturbations are inconsistent^14,25–27^, and (3) *spontaneous recovery* ^28–31^, the reversion back to previously adapted states due to removal of error or the passage of time. We decomposed adaptation into parallel implicit and explicit processes, and investigated if common experimental conditions facilitated or prevent their development and expression: (1) the effect of prior knowledge on explicit strategy^12,16,32^, (2) the ability to eliminate implicit learning by delaying error feedback^33–35^, (3) the ability to suppress strategic corrections by limiting the time individuals have to plan movement^12,35–38^, and (4) the ability to impair learning by increasing perturbation variability^35,39–41^.

Overall, we observed a reproduction of the in-lab results with the at-home experiments. We hope that the testing software provides researchers with the ability to rapidly and efficiently create at-home experiments solely through text file adjustments without low-level programming, thus allowing for longitudinal or cross-sectional studies of implicit and explicit learning in healthy and disease states.

## Results

We produced software that allows for at-home experiments and then used it to quantify sensorimotor adaptation along with its implicit and explicit components. We asked whether experimental conditions used to isolate these systems in the lab were effective in the uncontrolled at-home testing environment. All experiments were completed at-home and involved using the hand to move a cursor towards one of 4 targets either with a trackpad or a mouse on the subject’s Windows or Mac electronic device (Fig. 1B).

### Experience-dependent changes in learning rate: savings and anterograde interference

Previous work has demonstrated that if a subject experiences two matching or opposing perturbations, their experience can accelerate^15,21–23,35,37,42^ or slow^14,25–27,43^ further adaptation. We tested subjects in paradigms known to evoke this “savings” or “interference”. In Experiment 1, participants (n=16) were presented with two matching 30° visuomotor rotations (rotation 1 and rotation 2), separated by an intervening washout period (Fig. 1C, top). Subject hand angles during the adaptation and washout periods are shown in Fig. 1C (bottom). To compare performance duration rotations 1 and 2, both periods are shown overlaid in Fig. 1D, with pre-rotation hand angles subtracted to remove any bias (last 3 baseline cycles or washout cycles). To measure savings, we calculated the change in hand angle following the onset of rotation 2 (Fig. 1D, Δangle, change in hand angle 3 cycles post and pre rotation onset) and the empirical learning rate via an exponential curve (Fig. 1D, Δrate). Both adaptation metrics increased during rotation 2, consistent with savings (Δangle: t(15)=2.55, p=0.022; Δrate: t(15)=3.61, p=0.003).

In Exp. 2 (n=18), we exposed participants to two rotation periods (Fig. 1E, top) but reversed the perturbation’s orientation during rotation 2. As in Exp. 1, change in learning rate across rotation exposures (Fig. 1E, bottom) could be observed by comparing each rotation period’s learning curve (Fig. 1F; note, the initial reach angle is subtracted away to remove any bias). To measure whether learning was slowed due to interference, we calculated both the early change in hand angle as in Exp. 1 (Fig. 1F, Δangle, change in hand angle 3 cycles post and pre rotation onset) and the overall learning rate via an exponential curve (Fig. 1F, Δrate). We found that both measures decreased during exposure to rotation 2, consistent with anterograde interference (Δangle: t(17)=2.19, p=0.043; Δrate: t(17)=2.20, p=0.042).

In sum, both savings and anterograde interference emerged in the at-home setting, raising the possibility that modulation in sensorimotor adaptation can be studied at-home.

### Spontaneous recovery and the multiple timescales of adaptation

In addition to savings and interference, motor learning exhibits a third property: spontaneous recovery^28,29,31,44–46^. Models suggest spontaneous recovery is consistent with the presence of multiple adaptive processes, some fast and others slow^13,30,31^. McDougle et al. (2015)^19^ asked whether these states were differentially supported by implicit (subconscious) and explicit (strategic) processes.

In Experiment 3 (n=16), participants were exposed to rotation 1, which abruptly switched to rotation 2 to rapidly return behavior to near baseline (Fig. 1G, top). Reversing the perturbation’s direction was successful in washing out the overall adapted hand angle, resulting in a small negative bias that was not statistically significant (Fig. 1G, rotation 2, last 2 cycles, t(15)=1.25, p=0.231).

While a near-zero hand angle might suggest washout of adaptation, another possibility is that it was achieved through counterbalancing of a fast negatively biased explicit strategy, and slow positively biased implicit state. In such a case, ceasing explicit aiming should cause subjects to “spontaneously” revert toward the initial adapted state that was retained in implicit memory. To test this, we instructed participants to move their hand directly to the target and removed their visual feedback (Fig. 1G, dark gray area at right). As in McDougle et al.^19^, though participants aimed straight to the target, their hand angle rebounded to a positive level, i.e., toward the previously adapted state (Fig. 1H, no feedback). Indeed, a repeated-measures ANOVA across the baseline, end of rotation 2 (last 2 cycles), and no feedback periods, was consistent with spontaneous recovery (F(2,30)=8.41, p=0.001, η_p_^2^=0.359) with post-hoc tests suggesting no difference between the baseline and counter-rotation hand angles (p=0.362), but an elevation in no feedback hand angles over both earlier periods (no FB vs. baseline: p=0.001; no FB vs. counter-rotation: p=0.006).

In summary, in the at-home experiments we found evidence for multiple adaptive processes with different learning timescales, resulting in spontaneous recovery. This recovery could be mapped onto implicit and explicit states, as demonstrated in the laboratory^19^.

### Experience alters the competition between implicit and explicit components of adaptation

As previously reported^12,47^, Exp. 3 illustrated that laptop-based visuomotor adaptation is supported by implicit and explicit learning systems. In Albert et al. (2022)^12^, we studied the interactions between these two systems in the laboratory. A dominant phenotype was competition: when explicit strategy increased, implicit learning was suppressed. Thus, here asked whether implicit and explicit learning systems also compete with one another in the at-home experiments.

To answer this question, in Experiment 4, we tested two participant groups. One group consisted of naïve participants (n=25), people who had never experienced a visuomotor rotation either at-home or in-lab. In a second cohort, we tested participants who had never done a laptop-based rotation study but had experienced a visuomotor rotation during an in-lab study (n=10, “experience” group). Note, however, that experienced individuals had not recently participated in such experiments, with more than 6 months between in-lab and at-home studies. We wondered whether experienced individuals might transfer explicit knowledge gained about rotations from their in-lab to the at-home setting.

To test this, we exposed both groups to the same 30° visuomotor rotation (naïve and experience groups shown in Figs. 2A and 2B). Consistent with their past, participants that had an in-lab experience exhibited greater adaptation within 60 rotation cycles (total adaptation shown in Fig. 2C; t(33)=2.74, p=0.01). To test how this improvement depended on implicit and explicit learning systems, we instructed subjects to aim directly to the target and removed visual feedback of the cursor (gray region in Figs. 2A and 2B). As is typical, adaptation exhibited a sharp decline with this instruction, revealing implicit learning^12,47–50^. Explicit strategy could be estimated indirectly by subtracting total adaptation and the implicit learning measure. As suspected, adaptation accelerated in the experienced participants due to increased explicit strategy (Fig. 2D, t(33)=4.01, p<0.001); that is, participants with prior in-lab exposure were able to transfer their knowledge to the at-home setting.

**Figure 2.**
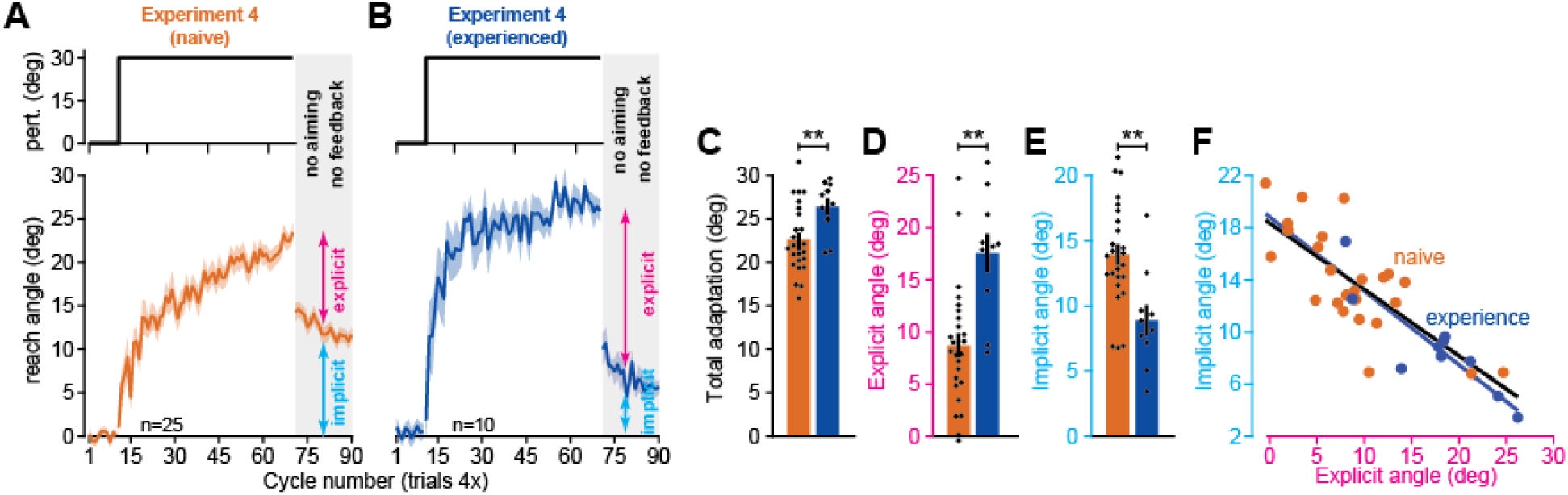
Experience increases strategy and inhibits implicit learning in the at-home setting. **A.** Naïve adaptation. Participants (n=25) were recruited that had not participated in an in-lab experiment before. The study started with a baseline period, rotation, and no feedback period during which subjects were told to aim straight to the target. Top: perturbation schedule. Bottom: movement angles. **B.** Same as in **A** except all subjects had previously participated (n=10) in an in-lab visuomotor rotation study (>6 months between this in-lab session and the subsequent at-home testing). **C.** Comparison between naïve and experienced total adaptation (mean angle over last 5 rotation epochs). **D.** Same as **C** but for explicit strategy (total adaptation minus implicit learning). **E.** Same as **D** but for implicit learning (mean angle over first 5 epochs in no feedback period). **F.** Regression of implicit learning onto explicit strategy (linear regressions fit separately to naïve and experienced cohorts). Error-bars indicate mean ± SEM. Points (**C**-**F**) indicate individual participants. Statistics in **C**-**E** denote unpaired t-test: **p<0.01.

This enhancement in strategy came with a cost: suppression of implicit learning. Though strategy increased, subconscious adaptation declined nearly 50% in experienced participants (Fig. 2E, t(33)=3.43, p=0.002). Our recent competition theory suggests^12^ that this inverse relationship is due to an error-based competition; when reliance on strategies is large, target errors are quickly reduced, depriving the implicit system of a critical learning substrate^51–53^. This theory is embodied by a competition model where implicit learning is proportional to the difference between the rotation’s size and explicit strategy, with a proportionality constant that depends on implicit learning properties; this is given by the linear equation *x_i_* = *p_i_*(*r* – *x_e_*), where *x_i_* is implicit learning, *x_e_* is explicit strategy, *r* is the rotation, and *p_i_* is the implicit learning gain (see Albert et al.^12^ and *Methods* for more details).

To test this competition model, we examined how implicit adaptation and explicit strategy varied across individual participants (Fig. 2F). Consistent with competition, linear regression suggested that the relationship between implicit learning and explicit strategy (lines in Fig. 2F) was similar across the naïve and experienced groups; regression slopes and intercepts differed by only 10.1% and 2.5%, respectively (slope 95% CI, naïve = [16.34,20.36], experience = [14.43,23.2]; intercept 95% CI, naïve = [-0.70°,-0.32°], experience = [-0.80°,-0.33°]). Furthermore, note that the competition model, *x_i_* = *p_i_*(*r* – *x_e_*), predicts that the regression slope (*p_i_*) and intercept (*p_i_r*) are linked to one another, differing only by the rotation’s size, *r*. Indeed, this also appeared consistent with the data, with the ratio between the intercept and rotation size (30°) differing from the regression slope by only 16.8% and 9.7% in the experienced and naïve groups respectively (0.51 vs. 0.61 in experienced; 0.57 vs. 0.63 in naïve group).

In summary, we observed that explicit knowledge acquired in the lab setting transferred to the at-home setting many months later, enhancing total adaptation. However, prior experience suppressed implicit learning via competition for error. This suggests that interactions between implicit and explicit learning systems can be studied in the at-home setting.

### Isolating the strategic learning system through delayed feedback

Implicit adaptation is thought to depend at least in part on cerebellar learning^9,54–61^, where plasticity at the Purkinje cell-parallel fiber synapses requires temporal correspondence between motor commands and error signals^62,63^. By extension, delaying error feedback should suppress implicit cerebellar learning. This idea has been tested in the laboratory numerous times, with clear reductions in implicit learning caused by a delay between the participant’s movement and the corresponding error^33–35^. In Experiment 5, we tested whether implicit adaptation could also be eliminated in a similar manner in the at-home setting, thus isolating strategic contributions to motor learning.

Subjects (n=21) moved the cursor to each target without visual feedback during the movement. The hidden cursor was rotated by 30°, frozen in place when it reached the target’s displacement, and revealed 1 second after the movement had completed. In Albert et al. (2021)^35^, we used similar conditions to eliminate implicit learning in the in-lab environment. In our at-home sample, we observed that 6 of the 21 subjects did not exhibit any adaptation (Fig. 3C). The notion that individuals could exhibit no adapted response at all was by itself is evidence for suppression of implicit learning, which exhibits an obligatory response to rotations^16,64,65^ even if participants do not intend to adapt.

**Figure 3.**
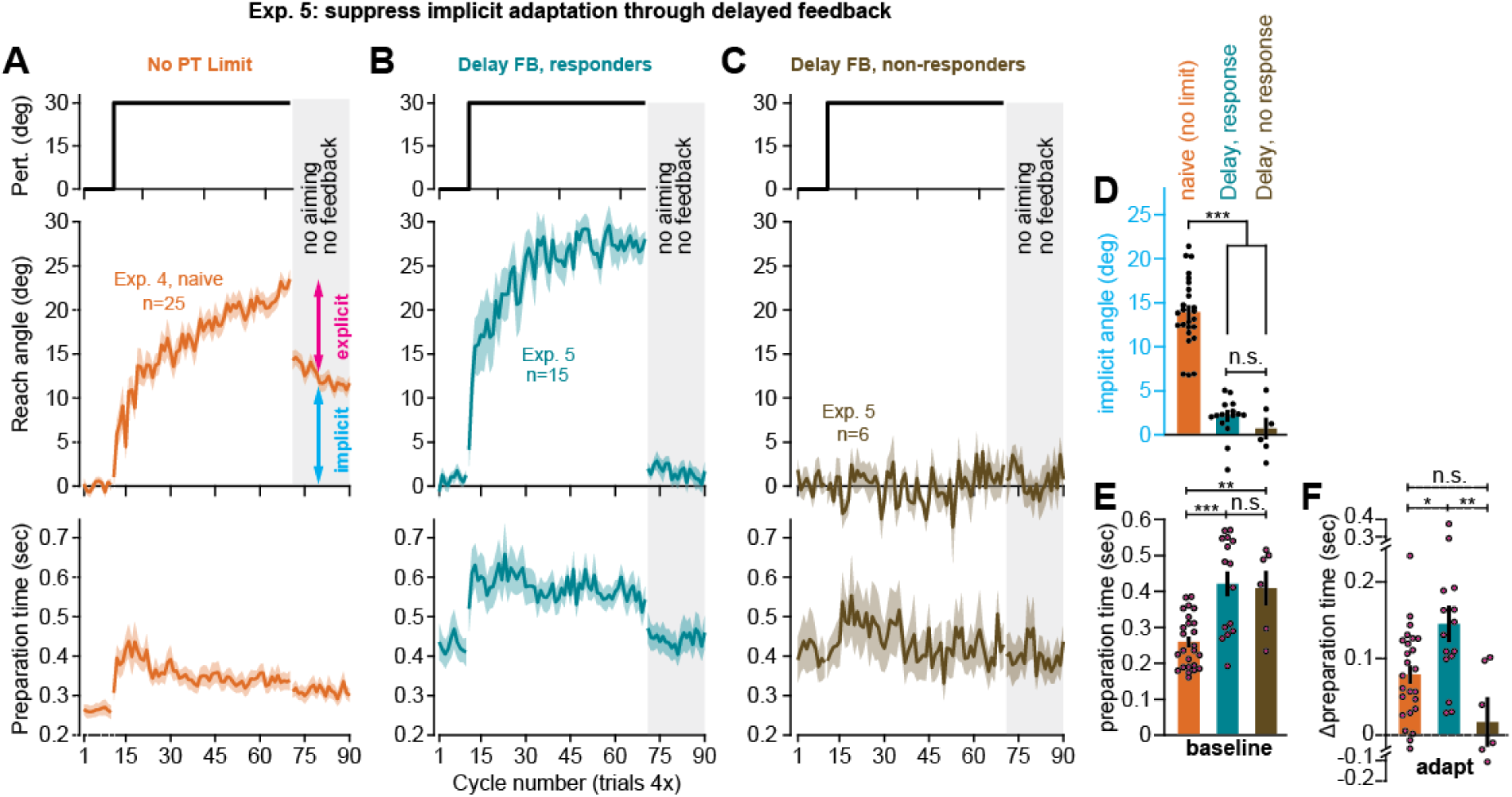
Delaying error feedback suppresses implicit learning in the at-home setting. **A**. Perturbation schedule, movement angles, and preparation time before movement onset for the naïve group in Experiment 4 (n=25). Preparation time is the latency between target appearance and movement initiation. **B.** Same as **A** but for participants that responded to the rotation (n=15) under delayed error feedback conditions (Experiment 5). **C.** Same as in **B** but for participants that did not adapt to the rotation (n=6) under delayed feedback conditions (Experiment 5). **D.** The implicit reach angle (average over first 5 no-feedback trials) for Experiment 4 and 5. Given small sample size in non-responder cohort, aftereffect was calculated over entire aftereffect period. **E.** Median preparation time during the baseline period (epochs 1-10). **F.** The change in preparation time during the rotation period (median movement angle over epochs 11-70, minus baseline preparation). Error-bars indicate mean ± SEM. Individual points (**J**-**L**) show individual participants. Statistics are one-way ANOVA: n.s. is not significant, *p<0.05, **p<0.01, ***p<0.001.

The remaining 15 subjects showed a brisk response to the rotation (Fig. 3B) with total adaptation exceeding that of naïve subjects who adapted with online feedback in Exp. 4 (compare Fig. 3A middle plot with 3B middle plot, t(38)=3.43, p=0.002). A similar increase in learning was observed in Albert et al. (2021)^35^, where residual error was eliminated in the delayed feedback group.

To test whether delayed feedback suppressed implicit learning, at the end of the rotation period we instructed subjects to move directly to the target without any cursor or error feedback, as in Exp. 4. We compared the implicit aftereffect during this no aiming period across the naïve participants in Exp. 4, as well as the responsive and unresponsive groups in Exp. 5 (Fig. 3D). Consistent with suppression of implicit learning, delaying feedback reduced the no aiming aftereffect (one-way ANOVA, F(2,43)=71.1, p<0.001, η^2^=0.768) by almost 90% in both the responsive group (post-hoc test against Exp. 4 control, p<0.001) and the unresponsive group (post-hoc test against control p<0.001). However, while the unresponsive cohort showed no statistically significant aftereffect (t(5)=0.58, p=0.588), the responsive participants in Exp. 5 showed a small aftereffect of approximately 2° (t(14)=3.3, p=0.005). This small bias is potentially consistent with reinforcement of successful movements during the rotation period^66^ and the creation of a use-dependent movement bias.

Finally, we considered how changes in implicit and explicit learning were reflected in movement preparation time. A well-known phenomenon in laboratory settings is that the use of explicit strategy requires preparation time, so that time-consuming mental rotations are completed prior to movement execution^18,37,38,67,68^. Indeed, we observed clear changes in preparation time in our at-home study (see Figs. 3E and 3F) across the control group (Exp. 4) and delayed feedback group (Exp. 5). For example, delaying feedback altered movement preparation time, irrespective of the rotation. Even during the baseline period, subject preparation time increased in delayed feedback participants (Fig. 3E, one-way ANOVA, F(2,43)=13.8, p<0.001, η^2^=0.391) with preparation time growing by approximately 60% in both the responsive (post-hoc test against Exp. 4 control, p<0.001) and unresponsive cohorts (post-hoc test against control, p=0.007). This overall change in movement preparation time is consistent with behavior in humans and other animals^69–73^ in response to variations in reward rate and the hyperbolic discounting of time; namely, delaying feedback increases the time to acquire the reward (i.e., feedback about hitting the target), and lengthens the inter-trial-interval, causing a reduction in response vigor^71,72^.

Given this initial bias in preparation time, to assess the relationship between explicit strategy and preparation time, the more critical quantity to measure is the change in movement preparation between the baseline and rotation phases^38^ (this subtraction removes any initial bias in preparation time unrelated to strategic re-aiming). This quantity mirrored explicit strategy use (Fig. 3F, one-way ANOVA, F(2,43)=7.18, p=0.002, η^2^=0.25), in that it was elevated only in the delayed feedback participants that responded to the rotation (post-hoc test against control, p=0.029; post-hoc test against unresponsive group, p=0.003), but not those who failed to develop an adapted response (post-hoc test against control, p=0.222).

In summary, we observed that implicit learning, explicit strategy, and movement preparation time were altered by delaying error feedback in the same manner across in-lab and at-home cohorts. Total implicit learning decreased approximately 90%. Explicit re-aiming increased and accelerated adaptation. Enhancement in strategy was accompanied by increases in movement preparation time.

### Isolating implicit learning by limiting reaction time

In Exp. 5, we found that an increase in explicit strategy is associated with longer movement preparation time^18,35,37,38,67^. By extension, in-lab studies have shown that restricting the participant’s reaction time reduces strategy use^12,18,35–38,68^. In Experiments 6 and 7, we tested this in the at-home setting (note that we reported a similar analysis in a previous study^12^).

Participants in Exp. 6 (n=21) and Exp. 7 (n=12) were tested under limited reaction time conditions (below, these groups are combined except where otherwise noted). On each trial, following target onset the participants were given a short reaction time window to begin their movement. Failing to meet the reaction time constraint resulted in a punishment tone and message at the end of the trial. This restriction was successful in reducing reaction time by approximately 120 ms relative to naïve control participants in Exp. 4 (compare Fig. 4A lower subplot with Fig. 4B and Fig. 4C, t(56)=5.17, p<0.001).

**Figure 4.**
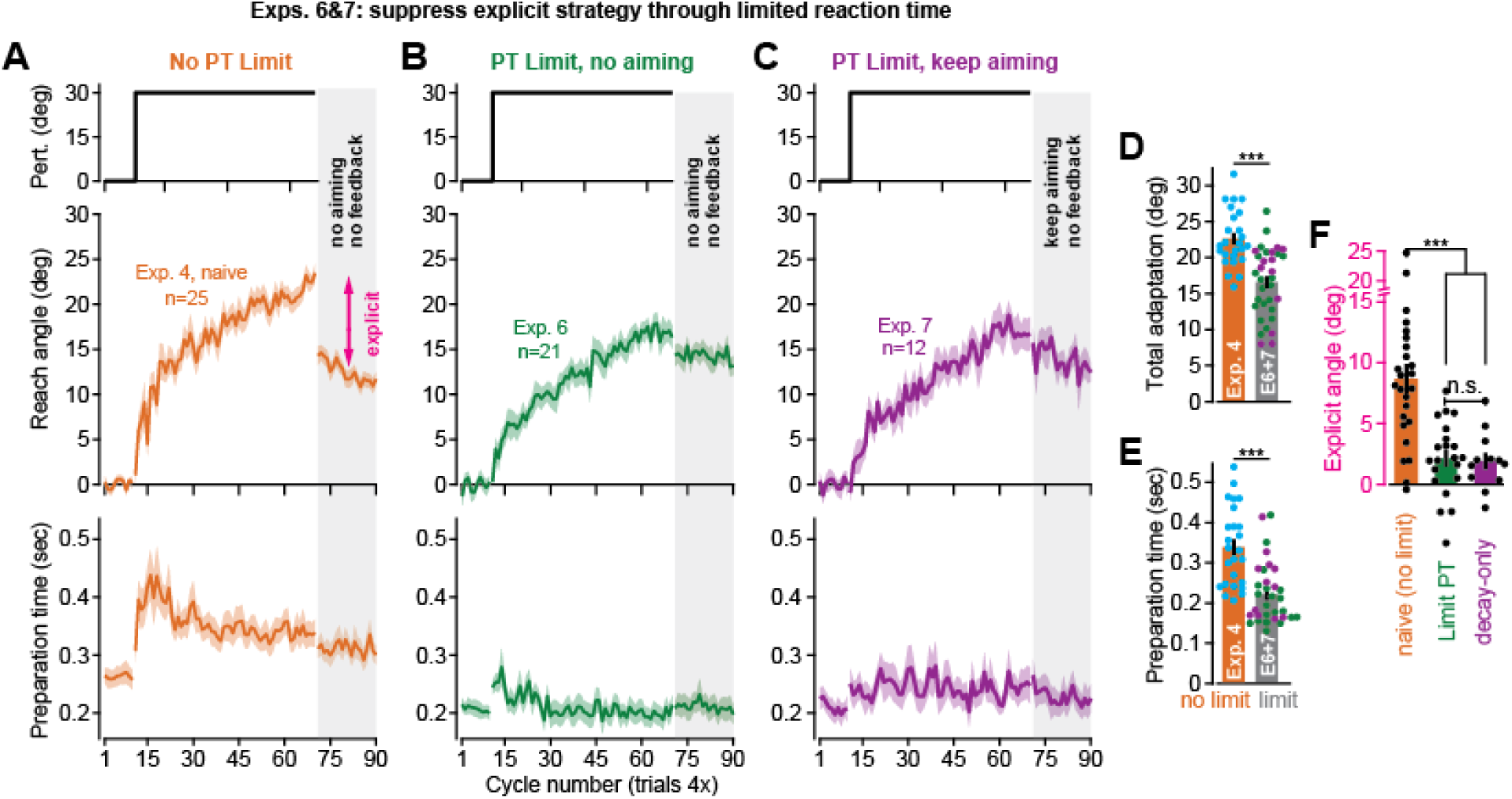
Restricting reaction time suppresses explicit strategy in the at-home setting. **A.** Perturbation schedule, subject movement angles, and movement preparation time for the naïve group in Experiment 4 (n=25). **B.** Same as **A** but for subjects (n=21) under constrained reaction time conditions in Experiment 6. Prior to the no-feedback period, participants were told to aim to the target and not to compensate for the previous rotation. **C.** Same as **B** but for subjects (n-12) under constrained reaction time conditions in Experiment 7. Prior to the no-feedback period, participants were told to continue to aim to move the imagined cursor to the target as they did in the perturbation period. **D.** Total adaptation over the last 5 rotation epochs. Experiments 6 and 7 combined because they were identical prior to the no-feedback period. **E.** Same as **D** but for preparation time during the rotation period (median over all 60 rotation epochs). **F.** Explicit strategy across Experiments 5-7 (total adaptation minus the implicit aftereffect: average angle over the first 5 no-feedback epochs). Error-bars indicate mean ± SEM. Individual points show individual participants. Statistics in **D** and **E** denote an unpaired t-test. Statistics in **F** denote a one-way ANOVA: n.s. is not significant, ***p<0.001.

We presented limited reaction time subjects with a 30° rotation. Consistent with suppression of explicit strategy, adaptation was slower and less complete with the imposed reaction time constraint (total adaptation shown in Fig. 4D; t(56)=5.13, p<0.001).

To test whether this change in total adaptation was due to a reduction in explicit strategy, subjects in Exp. 6 were instructed to aim their movement straight to the target, and all cursor and error feedback was removed (Fig. 4B, gray area at right). Explicit strategy was indirectly estimated by subtracting the hand angle during this aftereffect period from the total adaptation measured prior to the instruction. With that said, while this change in hand angle is primarily driven by participants abandoning their explicit strategy, this measure can be tainted by decay in temporally-labile components of implicit learning^30,50,74,75^. That is, approximately 30 seconds elapsed when participants received instructions to move straight without visual feedback, over which time some implicit decay is expected. To control for this decay, participants in Exp. 7 received instructions over a 30 second period but were not told to stop using explicit strategy (this group was told to try and hit the target with the imagined cursor as they had prior to the instruction period).

The changes in hand angle (post-instruction minus pre-instruction) are shown in Fig. 4E for Exp. 4 (control, no reaction time limit), Exp. 6 (decay due to strategy and time passage), and Exp. 7 (decay due to time passage alone). Critically, the instruction-driven change in hand angles was reduced by about 80% in both limited preparation time groups (Fig. 4L, one-way ANOVA, F(2,55)=16.1, p<0.001, η_p_^2^=0.369; post-hoc tests between control and Exp. 6 and Exp. 7, p<0.001). While the instruction-driven change in hand angle remained statistically significant in both Exp. 6 (t(20)=3.16, p=0.005) and Exp. 7 (t(11)=2.46, p=0.03), no statistically significant difference between these two groups was observed (post-hoc test, p=0.989). Thus, the small decrease in hand angle observed following instruction was likely unrelated to the cessation of explicit strategy, but instead simply the passage of time.

In summary, limited reaction time conditions provided a consistent way to restrict use of strategy both in-lab and at-home, providing a way to remotely track longitudinal changes in implicit learning.

### Altering motor adaptation through perturbation variability

In our last experiment, we tested another hallmark of sensorimotor adaptation: perturbation variability impairs motor learning^35,40,41,76^. As in our past in-lab experiments^35^, participants (n=26) in Exp. 8, were exposed to a rotation that varied trial-to-trial according to a normal distribution with a mean of 30° and standard deviation of 12° (Fig. 5A). Otherwise, the experiment was identical to the control group in Exp. 4 that experienced an invariant 30° rotation. Thus, the high-variance group in Exp. 8 had the same mean driving force (30°) as the zero-variance group in Exp. 4. Despite this, variability negatively impacted motor adaptation and led to slower and less complete learning (Fig. 5A, left and right column, last 5 cycles, t(49)=2.38, p=0.021). To determine whether this impaired response depended on implicit learning and explicit strategy, we instructed participants to move straight to the target without visual cursor or error feedback (Fig. 5A, gray area at right). As in Exp. 4, we interpreted the aftereffect during this period as implicit adaptation, and the difference between total learning and the aftereffect as explicit strategy.

**Figure 5.**
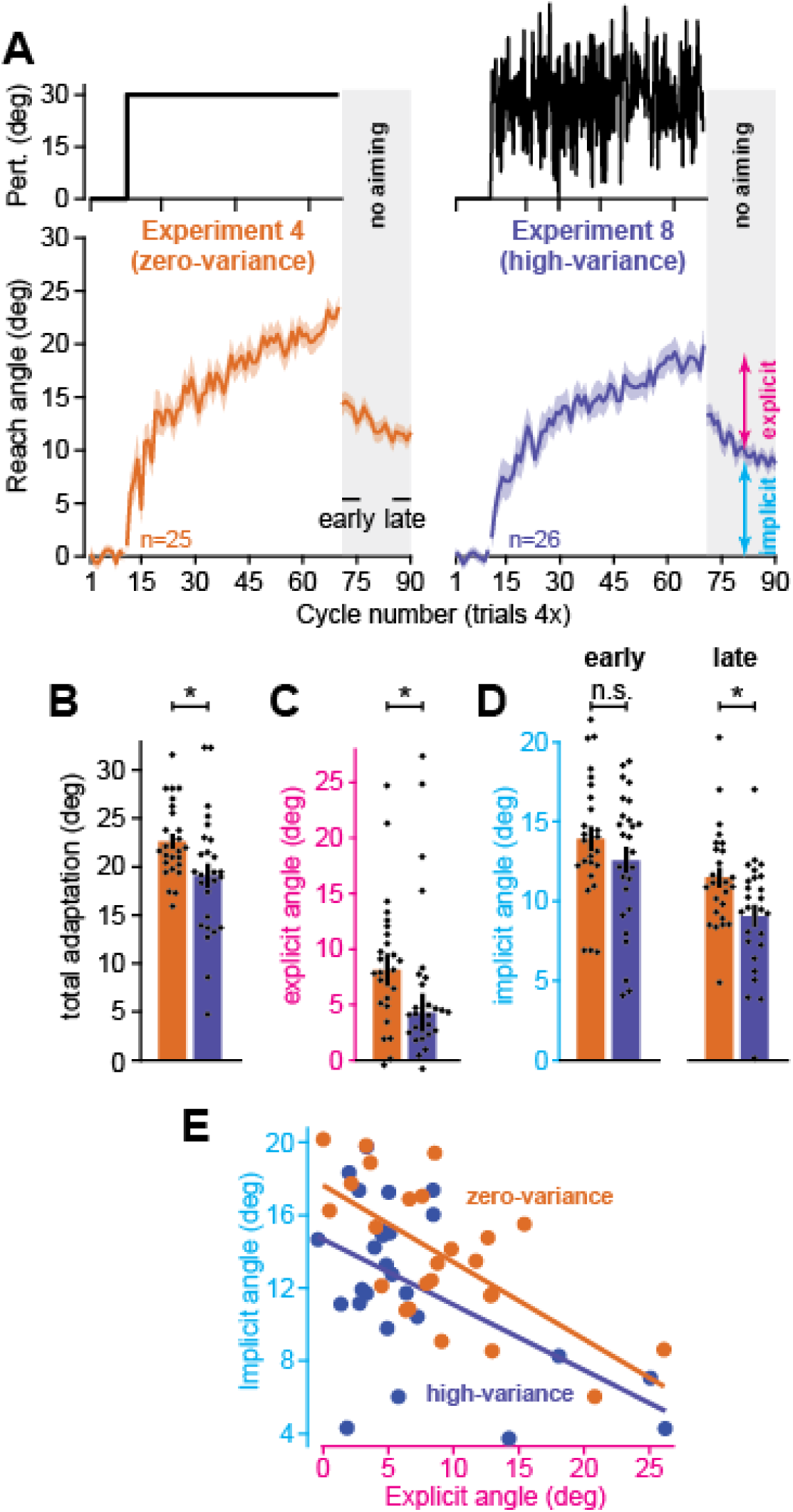
Perturbation variance suppresses implicit learning and explicit strategy. **A.** Left: perturbation schedule and movement angles for the naïve group in Experiment 4 (n=25). Right: same as left, but for participants in the high-variance group (rotations were sampled on each trial from a normal distribution (mean=30°, std. dev.=12°)). **B.** Total adaptation (last 5 rotation epochs). **C.** Explicit strategy (total adaptation minus implicit learning). **D.** Implicit learning: left is an early measure (first 5 no-feedback epochs), right is a late measure (last 5 no-feedback epochs). **E.** Uncorrelated measures for implicit and explicit adaptation. Because explicit equals total adaptation minus implicit learning, motor noise on implicit learning trials will be shared across both measures. To de-correlate them we took implicit and explicit measures using different trials: implicit (epochs 2, 4, and 6 of no-feedback period), and explicit (total adaptation minus epochs 1, 3, and 5 of no-feedback period). Solid lines show linear regression trendlines for each group. Error-bars indicate mean ± SEM. Individual points show individual participants. Statistics in **B** and **C** denote an unpaired t-test. Statistics in **D** denote Wilcoxon rank sum test. Data in **E** were entered into an ANCOVA: n.s. is not significant, *p<0.05.

While explicit strategies were highly variable across participants (Fig. 5C), perturbation variability suppressed the median explicit response to the rotation in the high-variance group (Fig. 5C, Wilcoxon rank-sum, z=2.2, p=0.028). Implicit adaptation, however, showed a more complicated pattern. First, the implicit response measured early during the no-aiming period appeared lower in the high-variance condition, but this trend was not statistically significant (Fig. 5D, t(49)=1.17, p=0.249). On the other hand, when we repeated this measurement at the end of the no aiming period (Fig. 5D, last 5 no-aiming cycles), high-variance implicit learning did show a statistically significant decline (t(49)=2.59, p=0.0123).

These implicit and explicit patterns differed from their in-lab analogues we observed in Albert et al. (2021)^35^. In-lab, we observed a decline in implicit learning alone, not explicit strategy. Secondly, in-lab differences in implicit learning were immediately evident on the first no-aiming cycle (they did not require additional movements to emerge, as in the at-home experiments).

We wondered whether these two observations were related. That is, while perturbation variance may have suppressed the implicit system’s response to error, decline in explicit strategy will increase the average error experienced by the implicit system, according to the competition equation: *x_i_* = *p_i_*(*r* – *x_e_*). Thus, high-variance participants in the at-home setting experienced two competing forces: implicit learning was impacted positively by an increase in error size (i.e., decrease in *x_e_* then increases *r* – *x_e_*), but negatively by a decrease in the implicit system’s responsive to that error (a decline in *p_i_*), each muting the other’s effect.

Altogether, changes in implicit learning cannot be understood independent of the explicit system. To this end, we considered the relationship between implicit learning and explicit strategy at the individual participant level (Fig. 5E) in an ANCOVA. Consistent with the competition theory, explicit strategy reduced implicit learning across both groups (ANCOVA, effect of explicit covariate, F=4.91, p=0.032, η_p_^2^=0.095). Now, controlling for explicit strategy (see *Methods*) revealed an impairment in the high-variance group’s early implicit learning measure (ANCOVA, fixed effect of variance, F=3.33, p=0.044, η_p_^2^=0.124).

In summary, introducing variance to the perturbation impaired adaptation in both the at-home and the in-lab settings. But this impairment mapped differently onto the implicit and explicit systems. In Exp. 8, explicit strategy exhibited a decline not observed in the laboratory. Implicit learning was impaired both in-lab and at-home, but the decline was stronger in the lab. The competition theory suggested the concomitant decline in explicit strategy partially masked the effect of variability on implicit learning in our at-home cohort.

## Discussion

Do experimental conditions known to enhance or suppress sensorimotor adaptation in the lab generalize to at-home studies? Here we examined this question across several diverse manipulations, and observed a correspondence between how humans behave in controlled laboratory settings and unmonitored at-home environments. The result is a tool that researchers can use to design and administer new experiments to examine implicit and explicit adaptation outside the laboratory.

### Experience-dependent changes in learning outside the laboratory

Learning processes change over time and with experience^35,39,42,77^. Understanding the conditions that can accelerate or hinder adaptation rates is a central question in motor control. A key feature of adaptation in both the cognitive^78^ and motor^23,79^ domain is savings: a facilitation in learning rate when a consistent perturbation is revisited. Many studies have documented this phenomenon in the laboratory ^23,31,77,79–81^. In Experiment 1 we examined this by presenting subjects with the same rotation separated by an intervening washout period. Participant behavior mirrored that described in-lab; the learning rate during the second exposure increased dramatically relative to the naïve state. On the other hand, reversing the second rotation’s direction during the re-adaptation period in Experiment 2 resulted in the opposite phenomenon: anterograde interference, i.e., the slowing of learning when two inconsistent perturbations are experienced in close temporal proximity^14,25–27,82^.

While both savings and interference were clearly detected in our at-home experiments, we did not make any attempt to localize them to implicit and explicit learning processes. There is a rich literature on the way that each component learning system contributes to savings, with many studies demonstrating a clear upregulation in strategic re-aiming^15,37^. Although we did not examine the issue, it is likely that this process occurred in our at-home experiment. For example, when we divided our cohort based on experience, we observed that individuals who had participated in a rotation study in-lab adapted much more rapidly to the same perturbation at-home, owing to a dramatic doubling in the use of strategy.

The role implicit learning plays in visuomotor rotation savings is currently debated, with various studies showing little to no increase^15,37,83^ in aftereffect during the re-adaptation period, and others^12,21,23^ pointing to a concomitant increase in subconscious motor corrections. In recent work^12^, we considered these apparent contradictions and proposed a competition theory which predicted that savings in the implicit system can be indirectly suppressed by large increases in strategic re-aiming (additional discussion below). While we did not investigate this idea further in the current study, competition between implicit and explicit processes was evident in other ways, most notably, in the substantial implicit learning decline exhibited by subjects with prior in-lab exposure to rotations.

In contrast to savings, few studies have examined the role that implicit and explicit systems play within anterograde interference. In a recent in-lab study, we limited reaction time to inhibit explicit aiming during an anterograde interference protocol, leading to a strong suppression in learning rate which could be attributed to the implicit system^12^. Though explicit learning has not been clearly assayed in interference protocols, some experimental paradigms^68,84^ with inconsistent rotations have been observed to facilitate subsequent re-adaptation, rather than degrade it. This hints at the possibility that explicit strategies may be less impacted by interference, and instead attempt to compensate for the deficit interference creates within the implicit learning system. The issue remains an open question.

All in all, much remains to be elucidated about the implicit and explicit mechanisms that support savings and interference, which we have now shown is possible to investigate in at-home studies. Follow-up studies could modify the paradigms described herein to measure implicit and explicit learning during training, and use preparation time constraints and error feedback delays to tease apart each system’s role by differentially suppressing the other.

### Multiple memory timescales outside the laboratory

The spontaneous recovery of memory is a fundamental behavioral property conserved across motor systems (e.g., saccades^28,44^, reaching^31,45,85^, vestibular reflexes^46^) and species^29^. Following adaptation, and subsequent washout, time passage causes movements to revert to a previously adapted state. Error-based learning models with multiple adaptive states can help to interpret this puzzling phenomenon^13,30,31^. For example, in a two-state leaning model, errors cause adaptation in two processes, one that responds strongly to error but poorly retains its memory over time, and another that learns weakly from error but exhibits robust retention. In this theory, previously adapted states are retained within the slowly-adapting process causing prior memory to reemerge when the rapidly adapting system dissipates.

Recent studies^19,86^ have examined this hypothesis as it pertains to visuomotor rotations and have proposed that implicit and explicit processes may differentially contribute to the slow and fast states that produce spontaneous recovery. Here we tested this phenotype in the at-home participants, finding that behavior mirrored that observed in the laboratory; though washout trials appeared to return subjects to their baseline sensorimotor state, the instruction to stop aiming caused the reemergence of a movement bias, due to a retained implicit memory. This experiment suggests that implicit learning has a similar persistence across in-lab and at-home environments, enduring over extended washout periods and thus producing spontaneous recovery.

It is important to stress that spontaneous recovery likely occurs via multiple mechanisms. That is, subconscious motor adaptation is likely supported by multiple adaptive processes that alone can produce recovery even without interplay with explicit strategy. For example, in force field adaptation, instructing participants not to compensate for the perturbation (told not push to prevent strategy use) still yields the standard spontaneous recovery timecourse^87^. In addition, subcortical areas that control saccadic^28,30^ and vestibulo-ocular adaptation^46^ also show a canonical spontaneous recovery time course. It remains an open question whether multiple implicit states also produce this type of spontaneous recovery in visuomotor rotation paradigms. To explore this, future studies could use our software to track implicit learning in a spontaneous recovery protocol, using reaction time constraints to minimize explicit re-aiming.

### Isolating implicit and explicit systems outside the laboratory

Laboratory-based studies have explored numerous tools to dissociate between the implicit and explicit processes hidden within a measured behavior. Here we examined two such manipulations: a delayed error feedback condition that reduces implicit learning^33–35^, and a limited reaction time condition that inhibits the ability to re-aim^12,18,35–38,68^.

Implicit learning relies at least in part on the cerebellum^9,54–61^, where LTD or LTP at Purkinje cell-parallel fiber synapses relies on the coincidence between simple spikes (efference copy of sensorimotor state) and complex spikes (error signal). Thus, delaying sensory error signals beyond the synapse’s plastic window should prevent subconscious motor adaptation in the cerebellar cortex^57^. This neurophysiological property appears consistent with behavioral phenotypes: when visual error is provided at a delay relative to movement, the implicit system’s learning is substantially attenuated^33–35^.

In Experiment 5, we observed that delaying feedback also strongly attenuated implicit learning in the at-home setting. While implicit learning was substantially reduced, it was not eliminated; there was a small, approximately 2° persistent bias in reach angle during the no aiming period. It is unclear whether this aftereffect was due to an error-based implicit learning process, or a use-dependent bias^66,88^ in reach direction that arose through reinforcement of successful actions during adaptation. In support of the latter, in the cohort of participants who showed no response to the rotation, this aftereffect was absent. With that said, sample size is small here, and should be examined in a larger follow-up study.

Explicit learning was also strongly altered by delaying error feedback. Whereas naïve participants with continuous feedback re-aimed by approximately 10°, subjects in the delayed feedback group (the cohort that responded to the rotation) more than doubled their aim magnitude (not shown in Fig. 3, total adaptation in Fig. 3B, middle plot, was approximately 27° with a 2° implicit aftereffect indicating 25° of reaiming). We also observed that explicit strategy was positively correlated with increases in movement preparation time. This is highly consistent with explicit phenotypes observed in the lab, with explicit strategy use increasing in response to feedback delays^33,35,67^, facilitated by increases in movement preparation time^18,35,38^. Interestingly, however, delayed feedback did not improve explicit strategy in all participants; some (6 out of 21) did not develop any strategy and exhibited no response to the rotation. This has also been observed in-lab, with a small subset of participants showing no re-aiming, across both short and long delays in error feedback^33^. Understanding why some individuals never develop an explicit strategy is a fascinating question that could be investigated at-home using our software package.

We explored a second manipulation designed to produce the opposite effect on explicit strategy: reaction time restriction. Several in-lab studies^18,38,68^ have shown that explicit planning is cognitively expensive, involving a mental rotation that can exceed naïve motor preparation time. Thus, by restricting the time individuals have to plan movement, explicit strategies can be tempered. In Exp. 6, we observed similar suppression in explicit strategy, in our at-home constrained reaction time task; nearly the entire adaptive response was mirrored within the implicit aftereffect trials, consistent with little to no re-aiming. To determine whether the small decrease in reach angle during the no-aiming period was due to residual explicit strategy or a temporal decay in implicit learning, we examined another participant cohort that was told to continue aiming during the no-feedback period. This group exhibited the same decline in reach angle, indicating that it was due to an unintended consequence of temporally labile implicit memory^50,74,89^ that decayed during the 30-second instruction period.

All in all, these preliminary data are encouraging that implicit systems can be isolated at-home, without having to instruct participants not to use an explicit strategy. With that said, the ability to isolate pure implicit learning via reaction time constraints can deteriorate over time, as the brain caches practiced explicit strategies and recalls them at short latency^18,68^. Follow-up studies are needed to better understand the experimental conditions that facilitate, or mitigate, explicit caching. For example, the total number of training targets (i.e., potential goals) is likely to correlate inversely with caching extent^18^. Further, it should be noted that our constrained reaction time task, which constrains preparation time *on every trial*, is quite different from in-lab conditions that limit reaction time on occasional trials. Can participants cache an explicit response in our task, without ever being given the chance to practice explicit re-aiming? These are critical questions that could be investigated at-home using our software package.

Thus, our results suggest that delayed error feedback and limited reaction time paradigms can be used at-home to differentially suppress implicit and explicit adaptation systems. We should also note that our software package includes an additional module commonly used to isolate implicit contributions to motor learning: invariant error-clamp^17,64,65^. This experimental manipulation has already been validated in a previous at-home study^10^, so we opted not to test its efficacy here (but has been included in our testing software for added utility, see *Methods*).

### Competition between implicit and explicit processes outside the laboratory

Uncorrelated trial-to-trial variability in the perturbation suppresses motor adaptation^12,39–41,76^. We recently observed that this decline was due to a disruption to the implicit system’s error sensitivity, decreasing its total response^35,39^. We found a similar impairment in implicit learning in response to perturbation variance in our online cohort, though the overall effect was considerably smaller: implicit learning dropped nearly 50% in our lab-based experiment, but only approximately 20% in our at-home experiment. Does this mean that implicit learning is more sensitive to variability within the in-lab context?

This might not be the case. Whereas explicit strategies appeared immune to variability within our lab-based study, our at-home cohort showed a sizeable decline (Fig. 5C, about 50%) in explicit aiming angle. This decline in explicit strategy could also modulate implicit learning, should these systems share at least one common error substrate^12,52^. In other words, suppose each system is engaged at least partly by visual error between the cursor and the target^12,51–53^. A smaller strategy will thus yield a larger target error to drive implicit learning. As a result, deterioration in implicit learning due to a reduction in its error sensitivity could be partly counteracted by an increase in the target error that drives implicit learning.

We recently studied this competitive (or compensatory) nature between the implicit and explicit adaptive systems across many in-lab studies^12^. This same competition rule appears to apply to implicit behavior in our at-home cohort. A key example is the comparison between naïve and experienced subjects in Experiment 4 (Fig. 2). Here, prior experience with a visuomotor rotation study in-lab caused marked enhancement in at-home adaptation, even though each testing period was separated at least 6 months in time. The culprit was explicit strategy, which subjects generalized based on their experience (Fig. 2D) much like upregulation in re-aiming observed with savings^15,68^. Remarkably, this improvement in strategy came at a substantial cost to implicit learning, which declined about 40% (Fig. 2E).

The competition theory clearly explains this otherwise puzzling result. Exposure to the visuomotor rotation paradigm long in the past creates an explicit memory. When the rotation was revisited, this past memory primed their explicit system, increasing the re-aiming magnitude. This priming increased the rate of adaptation (Fig. 2B), rapidly reducing error. But because the implicit system requires error to learn, this inadvertently suppressed the implicit response. In this competitive framework, experience does not change the inverse relationship between implicit and explicit learning (Fig. 2F); instead, it shifts where this antagonistic curve is sampled (naïve and experience in Fig. 2F are described precisely by the same linear regression but are translated along the regression line due to an increase in explicit strategy).

There are other sources that can contribute to negative covariations between implicit and explicit learning systems. First, consider the reverse to the scenario described above: explicit strategy may change in response to variations in implicit adaptation. This phenomenon has also been observed in the literature, with explicit responses declining to prevent implicit-induced rotation overcompensation^20,32,90^. Similarly, explicit strategy may increase in anterograde protocols (see discussion above), due to suppression in the implicit learning rate^12,68,84^.

However, there are ways in which implicit and explicit systems can *appear* to negatively co-vary, but only due to the way they are measured. Past studies have observed that implicit learning appears to generalize relative to aiming direction^48,86^. By extension, implicit aftereffect will decline when participants are told to aim in a direction inconsistent with their aim. Thus, an individual who aims further from the target, may appear to have less implicit learning^12,86,89^ when told to aim straight to the target during an aftereffect trial. We think these generalization curves will be critical to measure in future studies (see our past work^12^ for a more extensive analysis and discussion on this matter). While we did not measure implicit generalization curves in our current study, we have included a generalization module in our software to facilitate this process in follow-up studies (see *Methods*).

In sum, we think the interplay between implicit and explicit systems is central to understanding their complex behavior and learning rules. The competitive relationship between implicit and explicit systems was remarkably similar in our at-home studies to that observed in person, raising the possibility that interactions between these parallel learning systems can be studied in more detail in large at-home cohorts, both in cross-sectional and longitudinal studies.

## Methods

Here we detail the experimental protocols and data analysis used in this study. In addition, we reference supplementary information that details setting up, designing, and executing experiments with our at-home testing software package.

### Participants

The participants (n = 165; age range: 18-67) were neurologically healthy. 73 participants identified as male, 90 as female, and 1 answered N/A. 151 participants self-identified as right-handed, and 14 as left-handed. 94 participants used a Windows device, and 71 used a Mac. Participants were encouraged to use a touchpad; 143 participants used a touchpad, and 22 used a mouse.

We examined whether (1) using a trackpad or mouse, (2) having a Mac or Windows computer, or (3) being right or left-handed altered task performance. To do this, we considered the movement angles and reaction times during the baseline period (initial 40 trials prior to rotation onset in each experiment). We did not observe statistically significant differences in either measure based on individual-level factors (see Table 1 for statistical results and more detailed explanation).

**Table 1.**
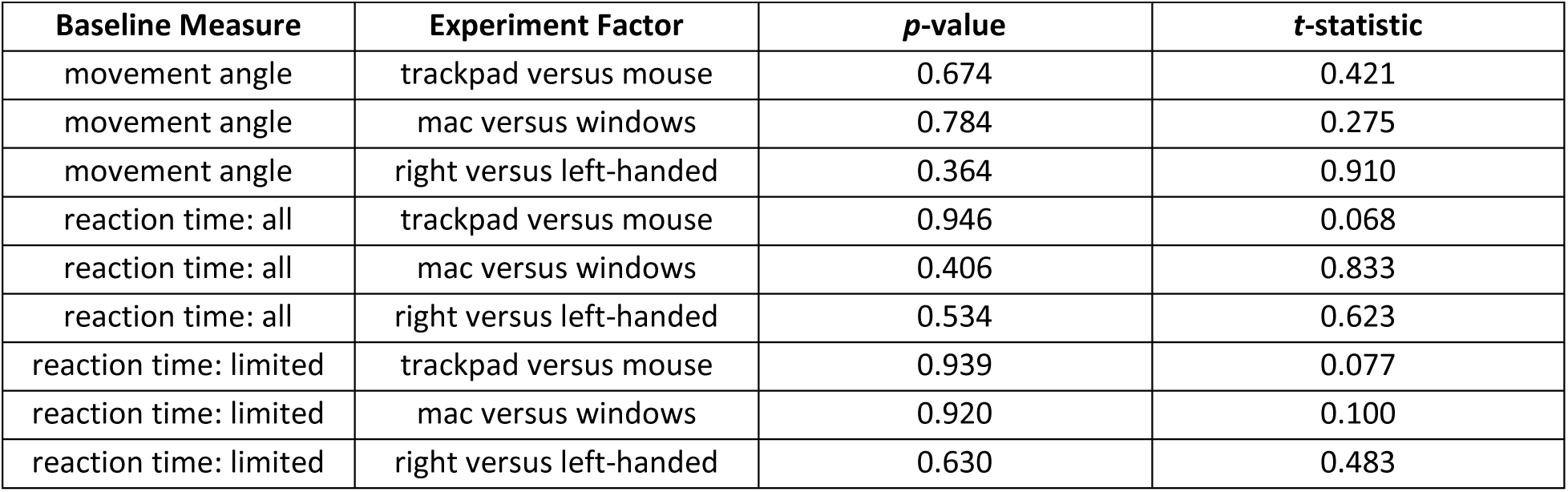
Movement angle and reaction time comparisons. We considered whether individual differences may alter movement parameters. Baseline Measure indicates what was tested: movement angle (average movement angle in the 40-trial baseline period, all experiments); reaction time: all (the reaction time during the 40-trial baseline period, all experiments); reaction time: limited (the reaction time during the 40-trial baseline period, but only experiments for which reaction times were not modulated by experimental manipulation, i.e., Experiments 5, 6, and 7 were not considered). Experiment Factor indicates the grouping variable tested: trackpad versus mouse (separated based on trackpad use or mouse use); mac versus windows (separated based on operating system); right versus left-handed (separated based on handedness). The *p*-values are uncorrected and correspond to two-sample t-test, with the test statistic provided in the *t*-statistic column.

| Baseline Measure | Experiment Factor | *p*-value | *t*-statistic |
| --- | --- | --- | --- |
| movement angle | trackpad versus mouse | 0.674 | 0.421 |
| movement angle | mac versus windows | 0.784 | 0.275 |
| movement angle | right versus left-handed | 0.364 | 0.910 |
| reaction time: all | trackpad versus mouse | 0.946 | 0.068 |
| reaction time: all | mac versus windows | 0.406 | 0.833 |
| reaction time: all | right versus left-handed | 0.534 | 0.623 |
| reaction time: limited | trackpad versus mouse | 0.939 | 0.077 |
| reaction time: limited | mac versus windows | 0.920 | 0.100 |
| reaction time: limited | right versus left-handed | 0.630 | 0.483 |

Subjects were recruited using several outreach tools: Facebook, Reddit, advertisements in group chats, posting on Johns Hopkins’ Announcements page, and word of mouth. Subjects were compensated at a rate of $10 per hour, in the form of Amazon gift cards delivered to the participant electronically. The protocol was approved by the Institutional Review Board at the Johns Hopkins School of Medicine.

### Apparatus

For all experiments, participants used their own computer or laptop in order to perform the task. The task was coded in Java; participants were instructed to start by downloading and installing Java onto their device. Then, participants were instructed to download and run a .jar executable. The executable contained a form to collect biographical data, an instruction video, a practice block of trials, and the experiment itself. The size and position of the experimental components (such as the cursor, target, and text prompts) were scaled based on the participant’s monitor size and resolution, automatically detected in software. After successful completion of the experiment, reaching data was saved as a .txt file, and participants were instructed by a window generated within the executable to upload this data file to a Dropbox file request. Data were recorded at a nominal sampling rate of 200 Hz (screen delay measurement is described below).

### Visuomotor rotation task

In all experiments, the subject was instructed to bring the cursor to the center starting position (red circle sized 0.015% of total screen size) at the start of each trial. After maintaining the cursor within this starting position for 1000 ms, a target circle (a green circle sized 0.04% of total screen size), appeared in 1 of 4 cardinal positions (0°, 90°, 180°, and 270°) at a radial displacement of 10 times the size of the target circle. The order of the four targets were randomized within each epoch (group of 4 trials). In addition, an audio cue (1000 Hz) was played to signal the start of the trial. Participants then performed a “shooting” movement by directing the cursor rapidly through the target without stopping. Once the target displacement was reached, the terminal position of the cursor was briefly ‘frozen’ in place.

At the end of the movement, subjects received auditory and visual feedback. If a participant reached too quickly (<10 ms) or too slowly (>275 ms), the target turned red or blue, respectively, and a ‘negative’ tone was played (200 Hz). For all paradigms, participants were also warned about their reaction time and movement speed when they were too slow (movement: >600 ms, reaction time: >1000 ms; these limits were more restrictive in limited reaction time conditions as described below) with text on-screen to reach or react faster. If a participant reached at the correct speed but missed the target, the target turned white, and a negative tone was played (200 Hz). If a participant reached at the correct speed and hit the target, the target turned purple, a positive tone was played (2000 Hz), and the participant received a point, which was tallied in the upper left hand corner of the experimental GUI. Participants were instructed to score as many points as possible. After feedback was given, the participant was instructed to move their hand back to the starting position to begin the next trial. The cursor remained hidden (but flashed occasionally every 500 ms for a duration of 200 ms per flash) until the cursor moved within 40% of the radial displacement.

Movements were performed in 3 possible conditions: null, rotation, and no feedback trials. On null trials, veridical feedback of cursor position was shown. On rotation trials, the cursor was rotated counterclockwise (CCW) relative to the start position. On no feedback trials, the cursor was hidden during the trial, and no audio or visual feedback was given regarding movement endpoint, accuracy, or timing.

To measure adaptation, we analyzed the movement angle on each trial. The movement angle was measured as the angle between the hand and the target (relative to the start position), at the moment where the hand exceeded 95% of the radial displacement between start position and target. This quantity was used exclusively for all quantitative analysis reported in this study.

### Preliminary instructions video

All experiments began with an instructional video, inserted within the executable file. The video showed a visual depiction of the testing environment and the trial structure, and played audio that highlighted the task details described in the previous section. In order to ensure that subjects paid attention to the video, a keyword briefly appears towards the end of the video; participants had to input this keyword in order to progress to the practice block, otherwise the instructional video was played again. Videos were tailored to 3 potential task environments: (1) the standard rotation task, (2) the limited reaction time task, (3) the delayed feedback task. The 3 videos are included online at: https://osf.io/e8b63/.

### Dynamic practice block

Following the instructional video, participants completed a practice block with null and no feedback trials. To progress to the experiment, the participant was required to complete successful movements on >65% of trials. A successful trial was determined if the subject reached and reacted satisfactorily quickly (<600 ms and <1000 ms, respectively), and reached within ±30 degrees of the target. When the desired accuracy was not achieved, the practice period was repeated.

### Software features, documentation, and code

Our at-home testing software consists of several modules that were examined in Experiments 1-8 (details provided below) supporting many experimental constructs: (1) constant abrupt perturbations, (2) variable abrupt perturbations, (3) no feedback trials, (4) delayed feedback trials, (5) limited movement preparation time trials, (6) implicit aftereffect trials, (7) savings paradigm, (8) anterograde interference paradigm, and (9) spontaneous recovery paradigm. Total number of targets and their locations can be modified and set breaks can be added at desired points during the testing protocol. Our software also has 3 modules that we did not test in at-home populations but were included given their common use in the literature: (1) invariant error-clamp trials^10,64,65^, (2) gradual rotation paradigm^12,91,92^, and (3) implicit generalization measurement protocol^48,86^.

The software package comes in two distributions. The first is a ‘lite’ standalone version that was designed to quickly and easily deploy new experiments. This version requires no programming proficiency and consists of an executable file plus a text file known as the ‘target file’. To make a new experiment, the code does not need to be accessed. Rather, the researcher can edit experiment parameters in the target file before running the executable. This process is thoroughly documented in *Supplementary File 1* and is also available in the *Instructions Manual* folder at: https://osf.io/e8b63/. This repository also contains the executable and target files described above that are used in generating user-defined experiments (in the *Code to Generate Experiments* folder).

Finally, experiments can also be generated through the ‘full’ software version. This distribution is designed for users that want to add or modify task features (e.g., experimental conditions, instruction sets, etc.) in our software package. A detailed instruction manual is located in *Supplementary File 2* (also https://osf.io/e8b63/), which describes the process of using and modifying the full software version (e.g., setting up the IDE, importing code, program layout, modifying code, generating executables, etc.).

### Measuring, total adaptation, implicit learning, and explicit strategy

In Experiments 3-8, we measured implicit learning during a specialized no feedback period. Prior to these aftereffect trials, the experiment paused for 30 seconds. During this time, a message was displayed on the screen, along with an audio track that instructed participants to move directly to the target during the no feedback trials without applying any method to compensate for the previous perturbation.

The average reach angle over the initial 5 epochs of this aftereffect period served as our implicit learning measure. To measure total adaptation, we averaged movement angles over the last 5 epochs of the rotation period. Explicit strategy was inferred by subtracting our implicit learning measure from our total adaptation measure.

### Screen delay measurement

We created a utility to measure temporal delay associated with updating the visual display on participant’s laptop computers. This utility (*Delay_Calculator.jar* available at https://osf.io/e8b63/) flashes a circle on the monitor and records participant button presses. The subject is told to press the button each time the circle flashes on the screen. Because the circle flashes at a regular interval, each circle flash can be predicted thus minimizing subject reaction time. Using the utility across several different machines and operating systems yielded an average delay of 154 ms. In all reaction time measurements reported here we adjusted these values by the average delay value. This utility was not used with all participants. Thus, we are using average delay value as an estimate for unmeasured screen delays. This is a potential source of error.

### Experiment 1: savings paradigm

This experiment was designed to test savings^15,21,23,37,77,81^ in motor output. Participants (n=16) completed the following trial structure (shown in Fig. 1C): 10 epochs of null trials, 30 rotation epochs with a 30° CCW rotation, 30 washout epochs (no rotation), and a 30-epoch exposure to the same 30° CCW rotation. Two set breaks were given (30 second rests) after trial 160 and trial 260.

Movement angles are shown over the training period in Fig. 1C. To compare the response to each rotation, movement angles during the second exposure period were sign reversed, and initial bias in reach angle was removed via baseline subtraction. The initial angle subtracted was calculated as the average angle over the last 3 epochs during either the baseline period (initial rotation) or the last 3 epochs during the washout period (second rotation). We quantified savings using an angular metric as well as a rate metric. The angular metric (Δangle) was calculated as the change in reach angle during the initial exposure to each rotation (mean angle over epochs 1-3 of each rotation, minus the preceding 3 epochs). We then calculated the difference in this angular change between the second and first rotation exposures. For adaptation rate, we fit a 3-parameter exponential curve to each learning curve that was constrained to begin at the movement angle on the epoch just prior to rotation onset. We next calculated the difference in the rate parameter for this exponential curve across each exposure (Δrate).

### Experiment 2: anterograde interference

This experiment was designed to examine anterograde interference^14,25,27,82^ in motor output. Participants (n=18) completed the following trial structure: 10 epochs of null trials, 30 rotation epochs with a 30° CCW rotation, 30 washout epochs, and a 30-epoch exposure to an opposing 30° CW rotation. Two set breaks were given (30 seconds rest) after trial 160 and trial 260. Baseline subtracted movement angles for each adaptation period were calculated as described above for Experiment 1. Interference was quantified using an angular measure (Δangle) and rate measure (Δrate) as described above for Experiment 1.

### Experiment 3: spontaneous recovery

This experiment was designed to detect spontaneous recovery of past motor memory^13,28,30,31^. Participants (n=16) completed a task in four blocks: 10 epochs of null trials, 60 epochs of perturbation 1 (abrupt, CCW 30°), 10 epochs of perturbation 2 (abrupt, CW 30°), then 20 epochs of no feedback trials. A single set break (30 seconds rest) was given after trial 160. To detect spontaneous recovery, we considered the movement angle over 3 critical periods: (1) baseline (cycles 1-10), (2) the end of the counter-rotation rotation (cycles 79 and 80), and the no-aiming period (cycles 81 to 100). Differences in movement angle across each period were analyzed using a repeated-measures ANOVA with post-hoc Tukey tests.

### Experiment 4: measuring implicit and explicit learning in naïve and experienced participants

This experiment assessed how experience with an in-lab visuomotor rotation protocol impacted learning in the at-home setting. We recruited naïve participants (n=25) who had no prior experience in a visuomotor rotation experiment in our laboratory. We recruited a second cohort (n=10) of participants who had completed an in-lab visuomotor rotation experiment in our laboratory. The trial structure was: 10 epochs of null trials, 60 epochs of a constant 30° rotation, 20 epochs of no feedback to measure the implicit aftereffect.

### Experiment 5: delayed error feedback

We tested whether delaying error feedback would suppress implicit adaptation^33–35,93^. Participants (n=21) completed a task with the same trial structure as Experiment 4. But unlike Experiment 4, we delayed visual feedback of the cursor position; subjects reached toward the target, but the cursor was hidden during the reach. Once the participant surpassed the radial displacement of the target, visual feedback of the cursor endpoint was provided after a 1055 ms delay.

Inspection of participant behavior suggested two groups in the data: subjects that adapted to the rotation, and others that were non-responsive (see Fig. 3-Supplement 1A for mean ± std. dev. of last 10 rotation epochs). Indeed, a Gaussian mixture model suggested that the data consisted of two components (AIC shown in Fig. 3-Supplement 1B). We used the model to sort subjects into each subgroup: ultimately, the model indicated n=15 individuals in the ‘responsive’ group and n=6 in the ‘unresponsive’ group. All analysis was conducted separately in each subgroup.

To determine whether implicit adaptation was suppressed by feedback delays, we calculated the movement angle in the no feedback period. For the responsive group, we calculated this angle as the mean movement angle over the initial 5 no feedback trials, as described above. Given the small sample size of the unresponsive cohort, we averaged over the entire no feedback period to minimize noise.

We next considered whether strategy use in each cohort was correlated to changes in movement preparation time. Because reaction times were prone to occasional extreme (large) values, we calculated the median reaction time over the baseline period (all 10 epochs) and the rotation period (all 60 cycles).

### Experiments 6 and 7: limiting reaction time

We tested whether limiting movement preparation time would suppress the participant’s ability to reaim^12,18,35–38^. Participants (n=33) completed a task with the same trial structure as Experiment 4. However, unlike Experiment 4, participants were required to start moving once the target had appeared. If their reaction time fell below a certain threshold on that trial, the participant was allowed to score a point for a successful movement. If their reaction time was above threshold, the participant was allowed to complete their movement and receive feedback, but no points could be scored and there was a ‘punishment’ message shown on-screen reminding them to react faster, along with a 200Hz tone.

The threshold that determined whether participants reacted satisfactorily or not was determined by an automatic calibration algorithm during the initial practice block. The practice period (80 null trials + 8 no feedback trials) was divided into two halves. During the first half of 44 trials, preparation times were not constrained. During the second half, a message appeared on-screen informing subjects that reaction times will be limited, and the threshold will be calibrated. The initial value of the threshold was set at 350 ms. During this phase of the practice block, the software measured reaction time within moving windows of 10 trials; if 8 or more trials had a ‘successful’ reaction time, the threshold decreased by 5 ms. Otherwise, the threshold increased by 5 ms. The threshold was capped between 280 ms and 350 ms: in other words, the participant could not calibrate their threshold outside this window. This procedure continued over the initial 40 baseline null trials during the experiment. The calibration period (84 trials total) terminated once the rotation was introduced and was maintained at the same level throughout testing. Note that this reaction time threshold was not adjusted for screen delay. Thus, given that screen delay was on average 154 ms, the true participant reaction time windows was bounded approximately between 126-196 ms.

This experiment included two separate groups. In Experiment 6 (n=21), participants were told not to use an aiming strategy and move their hand straight to the target during the final no feedback period.

In Experiment 7 (n=12), participants were told to keep aiming as if the perturbation were present, to move the imagined cursor through the target. Because these groups were identical up until the instruction, we combined them when comparing total adaptation and preparation time (median preparation time across the 60 rotation epochs) in Experiment 4. However, these groups were separated when comparing against Experiment 4 explicit strategy. We tested for differences between each group’s explicit strategy with a one-way ANOVA and post-hoc Tukey tests. Explicit strategy and total adaptation were calculated as reported above.

### Experiment 8: high-variance perturbation

In the last experiment, we tested whether perturbation variability impaired motor adaptation^35,40,41,76^. The protocol was identical to Experiment 4, except variability was added to the rotation. We used the same variance scheme as our in-lab study^35^: the rotation was sampled independently on each trial from a normal distribution (mean=30°, std. dev.=12°).

In addition to the ‘early’ implicit learning measure (average over initial 5 epochs), we also calculated the implicit aftereffect over the last 5 epochs. Note that we tested for differences in each learning measure using a one-sample t-test, with the exception of explicit strategy. This is because the explicit strategy in the high-variance group appeared to have outlying participants with particularly large strategies; accordingly, this group’s measure failed normality testing (Shapiro-Wilk test, W=0.733, p<0.001). Thus, we tested for differences in median strategies across the zero-variance and high-variance groups using a Wilcoxon rank sum test.

To explore how variance altered implicit learning while controlling for the adversarial relationship between implicit and explicit learning, we conducted an ANCOVA: implicit learning was the dependent variable, explicit strategy was the covariate, and variance-level was the fixed factor. However, the way we calculated explicit learning was dependent on implicit learning (see above). To avoid shared noise sources from inflating statistical measures in the ANCOVA, we recalculated implicit learning and explicit strategy with a different method. To calculate explicit strategy, we subtracted the average reach angle on the 1^st^, 3^rd^, and 5^th^ no feedback epoch from total adaptation. To calculate implicit adaptation, we averaged the movement angle on the 2^nd^, 4^th^, and 6^th^ no feedback epoch. Now, no shared epochs were used to calculate the implicit and explicit learning measures, ensuring no sharing of motor noise. We used this same process for the zero-variance (Experiment 4) and high-variance (Experiment 8) groups.

### Competition theory

Implicit and explicit adaptative systems are known to respond, at least in part, to target errors^12,51–53^. This means that these systems will necessarily exhibit competition. That is, when strategies are large, they will siphon target errors away from the implicit system, reducing its adaptation. We recently described this theory and its empirical evidence^12^. This theory is based on a state-space model where implicit processes adapt to target errors with error sensitivity, *b_i_*, and retain memory with a retention factor, *a_i_*. With long exposures to the rotation, the implicit system will eventually reach a steady-state: the point at which trial-to-trial error-based learning is counterbalanced by trial-to-trial forgetting. The competition model describes this steady-state according to: *x_i_*^ss^ = *p_i_*(*r* – *x_e_*^ss^). Here *x_i_*^ss^ represents steady-state implicit learning and *x_e_*^ss^ represents steady-state explicit strategy. The variable *r* is the rotation magnitude, and *p_i_* is an implicit learning gain determined by the parameters above: *p_i_* = *b_i_*(1 – *a_i_* + *b_i_*)^-1^.

This theory states that implicit adaptation will be proportional to the quantity *r* – *x_e_*^ss^ which is the amount of target error that is left when the participant reaches their steady-state explicit strategy. Thus, as strategies increase, this residual target error will decrease, then reducing implicit learning. We use this model to interpret implicit behavior in Experiments 4 and 8 (Figs. 2F and 2G). For a more comprehensive analysis and discussion on these matters, see our recent work^12^.

### Data and code availability

The data and analysis code will be made available upon publication.

## Acknowledgements

This work was supported by grants from the National Institutes of Health (R01NS078311, F32NS095706), and the National Science Foundation (CNS-1714623).

---

**Figure 3-Supplement 1.**
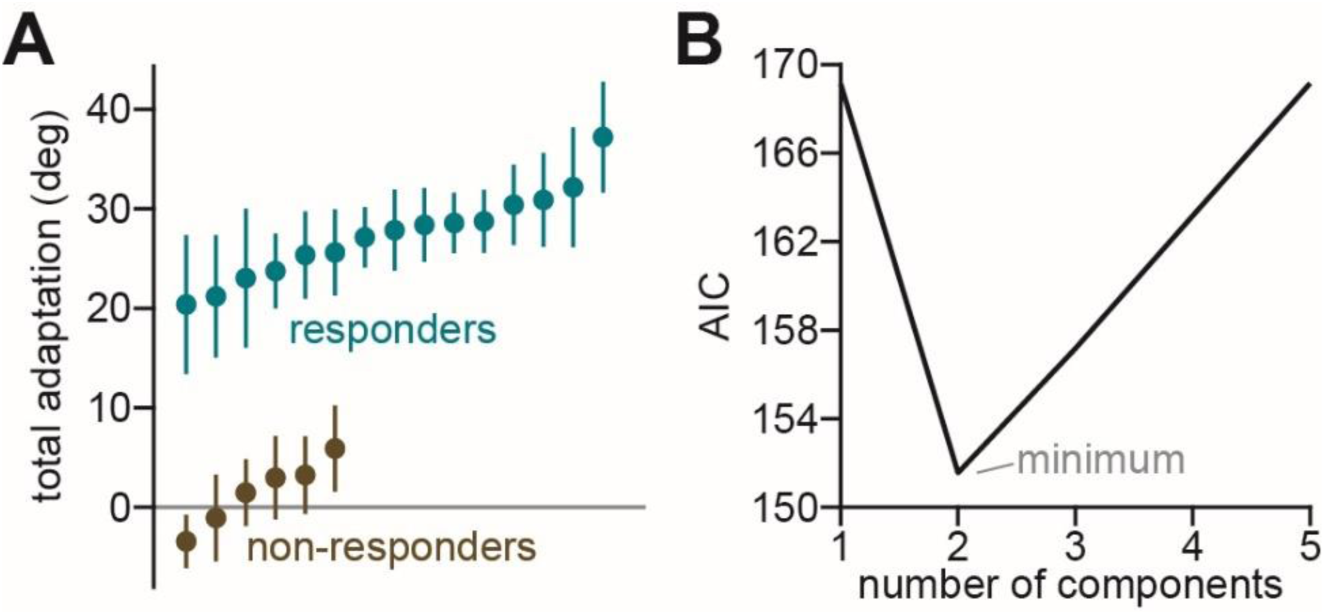
Two subgroups in the response to delayed error feedback. **A.** We calculated the movement angle over the last 10 rotation epochs in Experiment 5 (n=21). Visually, two subgroups were apparent in the data. **B.** To rigorously test whether participants fell into two groups, we fit a Gaussian mixture model, and varied the number of components between 1 and 5. We selected the model with the smallest AIC value (2 components).

