## Supplementary File 1 for "A software tool for at-home measurement of sensorimotor adaptation"

### Introduction

This software will allow you to run visuomotor rotation experiments using a laptop or PC. Subjects participate using a touchpad or mouse in order to make "reaching" movements with the mouse cursor. The data is outputted as a text file, and can then be parsed and analyzed using the language of your choice.

### What Experiments Can I Run?

This standalone version is able to support the following experiment types/parameters:

- Abrupt perturbations, with or without variance
- Gradual perturbations
- No feedback trials
- Error clamp "perturbation" trials
- Delayed feedback paradigm
- Limit reaction/preparation time paradigm (with automatic reaction time threshold calibration)
- No feedback implicit probe paradigm
- Generalization paradigm
- Savings paradigm
- Interference paradigm
- Spontaneous recovery paradigm
- Multiple targets
- All experiments can have set breaks

### Requirements

Currently, this software only runs on Windows and Mac. The device must have Java installed: https://java.com/en/download/

We also recommend that subjects participate using a device that has the capability of playing audio, so that they are able to hear the sounds played during the experiment.

### How To Get Started

TargetFile.txt must be placed in the same folder as the Standalone.jar file. After making sure that the parameter values inside TargetFile.txt are correct, simply double click Standalone.jar. The program will automatically start up using the parameters specified in the target file. If parameter values conflict (i.e. you set parameters saying that the experiment is both a savings task and an interference task), then the experiment will not run. You can run the .jar in your command line using the following command:

```
java -jar Standalone.jar
```

in order to see the errors that are being outputted.

Also, be sure to type in all values in the target file in the following format:

```
[field name]: [field value]
```

Example:

```
num_baseline: 40
```

Finally, do not delete the line of six equal signs (======); this line of characters tells the code where to delineate between the comment at the top of the text file and the parameter values below.

It is up to you how this experiment is conducted. You could have a device in your lab space that runs this software, and subjects could come to you directly to participate on this device. Alternatively, you could send out the .jar and .txt files to participants via the internet, and have them participate remotely and send back the data files that they generate.

Note that this version of the software only encompasses the experiment itself - it does not collect biographical data, play instructions, include a practice round, etc. You could potentially have the subject access a website that collects biographical data and plays an instructional video before directing them to a download link for the experiment.

For practice, you could create two different standalone versions - one for practice and one for experimentation - and have the subject complete the two tasks back to back. Alternatively, you could include a very long baseline period for practice, and then add a set break in order to differentiate between practice and experiment.

Finally, you could access the full version, where you can download the Java code directly and generate an experiment with a built-in practice block, instructions video, etc. More information here:
https://docs.google.com/document/d/1oL1jncx91ZkzWAWWmNVcqxKZGkFbbcGrYY54dSeJNLM/edit?usp=sharing

### What Do the Parameters Mean?

All values should be integers unless otherwise stated.

- **num_baseline** - Specifies the number of baseline (null) trials in the experiment.
- **num_rotation** - Specifies the number of perturbation trials in the first half of the experiment. If not running savings, interference, or spontaneous recovery, then the number of trials indicated here is the total number of perturbation trials in the experiment.
- **num_washout** - Specifies the number of washout trials in the experiment. Set this value to 0 if not running savings, interference, or spontaneous recovery.
- **num_rotation_2** - Specifies the number of perturbation trials in the second half of the experiment, if running saving/interference/spontaneous recovery.
- **num_no_feedback** - Specifies the number of no feedback trials at the end of the task.

Note: the experiment assumes that the order of trials for non-generalization experiments is baseline -> rotation -> washout -> rotation_2 -> no_feedback. If you wanted to, for example, run a simple abrupt perturbation experiment with no feedback trials at the end, then just set num_washout and num_rotation_2 equal to 0. You can't, however, have washout occur after rotation_2.

- **num_generalization_rotation** - Specifies the number of perturbation trials that are present in each cycle of generalization.
- **num_generalization_NF** - Specifies the number of no feedback probes during each cycle of generalization.
- **num_generalization_cycles** - Specifies the number of generalization cycles.
- **angle_range_1** - Specifies the lower bound angle for generalization no feedback probes. This value should be a double (i.e. has a decimal point), and is in degrees.
- **angle_range_2** - Specifies the upper bound angle for generalization no feedback probes. This value should be a double (i.e. has a decimal point), and is in degrees.

Note: for generalization tasks, the structure of trials is num_baseline -> num_rotation -> (num_generalization_rotation + num_generalization_NF)*num_generalization_cycles. During the rotation portion of the cycle, the task is identical to the perturbation block - subjects will reach towards targets with a standard VMR perturbation. During the no feedback portion of the cycle - for every no feedback probe trial, a target will randomly appear at evenly spaced locations between angle_range_1 and angle_range_2. (For example, if we set num_generalization_NF = 3, angle_range_1 = 70, and angle_range_2 = 90, then during this portion of the cycle, we would have three reaches to three targets: one appearing at 70 degrees, one at 80, and one at 90.)

- **num_targets** - Number of targets in the experiment, evenly spaced. The order of the targets are randomized during the experiment.
- **ITI** - The amount of time that the subject must spend on the start dot before the trial starts. This value should be a double (i.e. has a decimal point), and is in ms.
- **minimum_reach_time** - The minimum amount of time that the subject must spend during their reach. If they reach faster than this, then they are penalized for moving too fast. Measured in ms.
- **maximum_reach_time** - The maximum amount of time that the subject must spend during their reach. If they reach slower than this, then they are penalized for moving too slow. Measured in ms.
- **warning_reach_time** - If they reach slower than even this, then they are penalized for moving too slow, and also receive a warning on the screen. Measured in ms.
- **non_limit_rt_threshold** - During a non limit reaction/preparation time trial, if a subject reacts to the trial slower than this threshold, then they are penalized for moving too slow, and also receive a warning on the screen. Measured in ms.
- **angle_mean -** Specifies the angle of perturbation during the num_rotation block. If running a gradual paradigm, this is the angle that the experiment plateaus at. A negative angle means the cursor is rotated counterclockwise. This value should be a double (i.e. has a decimal point), and is in degrees.
- **angle_std** - Specifies the standard deviation of the perturbation. If this value is non zero, the experiment will randomly generate perturbations from a normal distribution using the programmed value for angle_mean as the mean, this value as the standard deviation. This value should be a double (i.e. has a decimal point), and is in degrees.
- **angle_mean_2 -** Specifies the angle of perturbation during the num_rotation_2 block. If running savings, make sure that angle_mean = angle_mean_2. This value should be a double (i.e. has a decimal point), and is in degrees.
- **angle_std_2** - See entry for "angle_std"
- **is_delayed_feedback -** Integer flag to indicate if this experiment should run a delayed feedback paradigm. Setting it to "1" means it is delayed feedback; otherwise, set it to 0.
- **delayed_feedback_delay** - The duration of the delay before feedback is shown during delayed feedback trials. Measured in ms.
- **is_limit_rt -** Integer flag to indicate if this experiment should run a limit reaction/preparation time paradigm. Setting it to "1" means it is limit RT; otherwise, set it to 0.
- **rt_threshold -** During a limit reaction time paradigm, if the subject reacts slower than this threshold, then they fail the trial and are "punished" with a warning message and long tone. This value is automatically calibrated during the baseline period of the experiment in order to ensure that subjects are able to perform the task with some success. The value you place here in the target is the initial starting point of calibration. Measured in ms.
- **angle_threshold -** During calibration, this threshold determines the angle at which the subject must move within in order for that calibration trial to be successful. For example, if this threshold is set at 30 degrees, but the subject reaches towards the target at 40 degrees (either clockwise or counterclockwise to the target), then the trial is counted a failure, even if the subject reacted quickly enough. This field should be a positive double, measured in degrees.
- **lower_rt_cap** - The lower bound for RT threshold calibration; rt_threshold can not go lower than this value. Measured in ms.
- **upper_rt_cap** - The upper bound for RT threshold calibration; rt_threshold can not go higher than this value. Measured in ms.
- **RT_calibration_window -** The length of the moving window that the calibration algorithm uses. Measured in number of trials (i.e. a value of 10 means that the algorithm looks at a moving window of 10 trials during baseline).
- **RT_calibration_threshold -** The threshold of success that must be attained within a moving window in order to decrease the rt_threshold value. Otherwise, this value will increase. Must be a double, and a value between 0 and 1.
- **RT_calibration_increment** - The amount in which the rt_threshold value is changed depending on the success or failure of the moving window. Measured in ms.

Note: The calibration algorithm works as follows:
During baseline, subjects will reach towards targets in a null condition. Within a moving window of trials defined by RT_calibration_window, each individual trial will be classified as either pass ("1") or fail ("0"). In order to pass, the reach must be within the threshold defined by angle_threshold, and the subject must react faster than the value currently assigned to rt_threshold. Then, the trials in the moving window are assessed to see if the percentage of successes is greater than or lower than the threshold defined by RT_calibration_threshold. If success is greater, then rt_threshold is lowered by a value equal to RT_calibration_increment, and vice versa. If a subject reaches either lower_rt_cap or upper_rt_cap, then rt_threshold cannot be incremented further above or below these caps, even if the moving window is successful or not, respectively.

- **is_no_feedback_implicit_probe -** Integer flag to indicate if this experiment should run a no feedback implicit probe paradigm. Turning this flag on means that before the no feedback block of trials, the experiment pauses and displays a message (with a voiceover) that instructs subjects to reach straight to the target. Setting it to "1" means it is an implicit probe paradigm; otherwise, set it to 0.
- **is_EC -** Integer flag to indicate if this experiment should run an error clamp paradigm, where reach movements during the rotation period are clamped at the angles specified by angle_mean and angle_mean_2. Setting it to "1" means it is an error clamp paradigm; otherwise, set it to 0.
- **is_gradual -** Integer flag to indicate if the perturbation is gradual, rather than abrupt. Setting it to "1" means it is a gradual paradigm; otherwise, set it to 0.
- **num_plateau -** Specifies the number of plateau trials that will be present after the perturbation is gradually ramped up to the maximum value specified by angle_mean.

Note: If running a gradual paradigm, the rotation block is constructed as follows: the total number of perturbation trials is encoded by num_rotation. Then, this block is divided into two parts: the gradual ramping up, and the plateau period. The number of plateau trials is specified by num_plateau; from there, num_gradual is simply num_rotation - num_plateau.

- **is_generalization -** Integer flag to indicate if this is a generalization paradigm. Setting it to "1" means it is a generalization paradigm; otherwise, set it to 0.
- **post_reach_delay -** Determines the amount of delay at the end of each trial. After feedback is given, the experiment momentarily freezes in order to give the subject time to process their performance. Measured in ms.
- **is_savings -** Integer flag to indicate if this is a savings paradigm. Setting it to "1" means it is a savings paradigm; otherwise, set it to 0.
- **is_interference -** Integer flag to indicate if this is an interference paradigm. Setting it to "1" means it is an interference paradigm; otherwise, set it to 0.
- **is_spontaneous_recovery -** Integer flag to indicate if this is a spontaneous recovery paradigm. Setting it to "1" means it is a spontaneous recovery paradigm; otherwise, set it to 0.

Note: In order to generate a spontaneous recovery paradigm, you can utilize the following structure: baseline -> rotation -> rotation_2 -> no feedback. Set num_washout = 0.

- **set_break_length -** The length of set breaks during the experiment. Measured in ms.
- **set_break_trial_nums** - Specifies the trials at which set breaks occur. For example, if trial number 5 is indicated here, then the set break happens between trial 4 and trial 5. For multiple set breaks, separate trial numbers using commas (e.g. 40, 120, 260)
