## Supplementary File 2 for "A software tool for at-home measurement of sensorimotor adaptation"

### Introduction

This software will allow you to run visuomotor rotation experiments using a laptop or PC. Subjects participate using a touchpad or mouse in order to make “reaching” movements with the mouse cursor. The data is outputted as a text file, and can then be parsed and analyzed using the language of your choice.

In the full version, you can generate full experiment files directly (without the use of a target file), so that subjects only need to download and run a single executable file in order to participate.

### What Experiments Can I Run?

This code is able to support the following experiment types/parameters:

- Abrupt perturbations, with or without variance
- Gradual perturbations
- No feedback trials
- Error clamp “perturbation” trials
- Delayed feedback paradigm
- Limit reaction/preparation time paradigm (with automatic reaction time threshold calibration)
- No feedback implicit probe paradigm
- Generalization paradigm
- Savings paradigm
- Interference paradigm
- Spontaneous recovery paradigm
- Multiple targets
- All experiments can have set breaks
- The full code is provided, so that you can add, remove, and edit any features you want so that you can create experiments that are uniquely yours

### What Else Is Included?

In addition, you can:

- Include a built-in consent form for biographical data collection
- Include a YouTube video with instructions
  - This video can be embedded within the Java GUI on Windows; otherwise for Mac, a hyperlink is provided instead
- A customizable practice block of trials
- A closing window with links to allow subjects to directly upload their data to a file hosting service (Dropbox, OneDrive, etc) of your choice
- The full code is provided, so that you can add, remove, and edit any features you want so that you can create experiments that are uniquely yours

### Requirements

Currently, this code only generates software for Windows and Mac. Each device that runs this software must have Java installed: https://java.com/en/download/

We also recommend using the Eclipse IDE in order to edit and compile this code on your machine: https://www.eclipse.org/downloads/packages/

However, any Java IDE of your choosing should work just as well.

Finally, we also recommend that subjects participate in these experiments using a device that has the capability of playing audio, so that they are able to hear the sounds played during the experiment.

### How to Use This Code

#### Setting up Java + IDE

First, download Java using the following link: https://java.com/en/download/ . Installation for Java should be fairly straightforward - just follow the instructions provided by the installer.

Next, Java code can be edited using any Java Integrated Development Environment (IDE), but we recommend Eclipse. First, download Eclipse and run the installer: https://www.eclipse.org/downloads/packages/ . Choose the first option on the menu that pops up, “Eclipse IDE for Java Developers”:

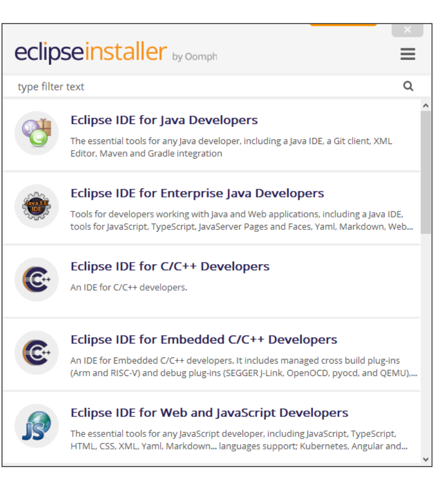


and proceed with the installation.

Once installed, run Eclipse. You will get a message asking to select a directory as your workspace - usually, the default option works fine. Your workspace is the directory in your file system that houses the code, resource files, etc that you will be working with. You can move this folder anywhere within your file system, and browse for it in this window before proceeding.

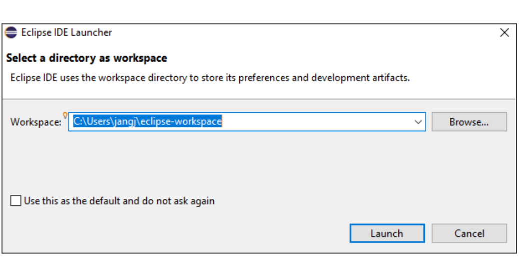


Next, you will see the following Welcome screen. Feel free to “X” out of it:

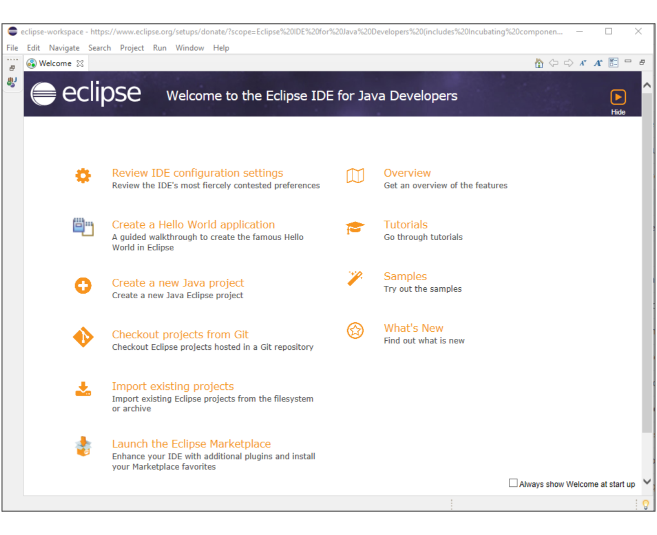


You will then see the main IDE GUI. Before continuing, we recommend adding a “Console” window to the GUI in order to make things easier for yourself in the future. The console will print out errors and print statements, which will help with any debugging that you need to perform. In order to add the “Console,” simply go to Window > Show View > Console. The Console will appear at the bottom of the GUI, and will look like this:

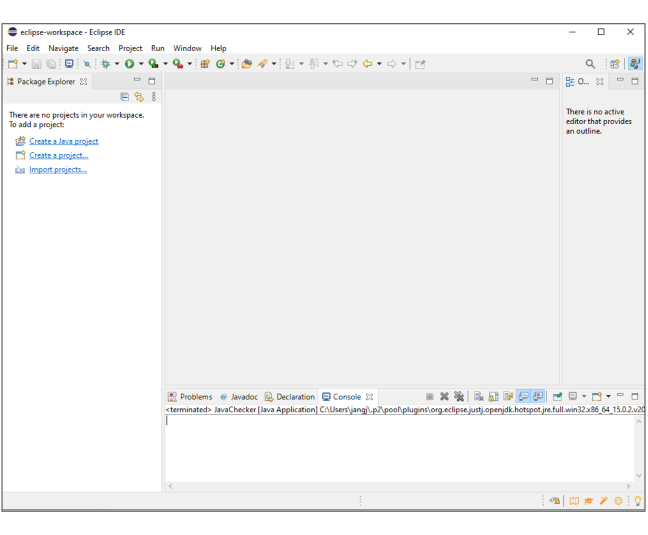

#### Importing the code

Download the code and its resources (.zip file). Then, in Eclipse, click File > Import > General > Existing Projects into Workspace, and click Next.

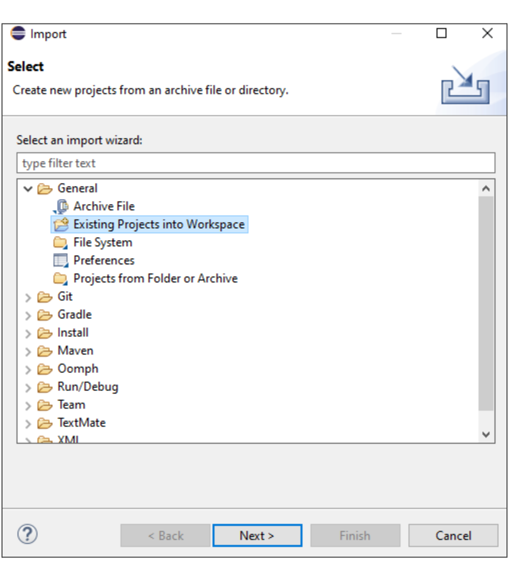


Next, click “Select archive file” and browse within your file system for the .zip file you just downloaded. Click “Finish.”

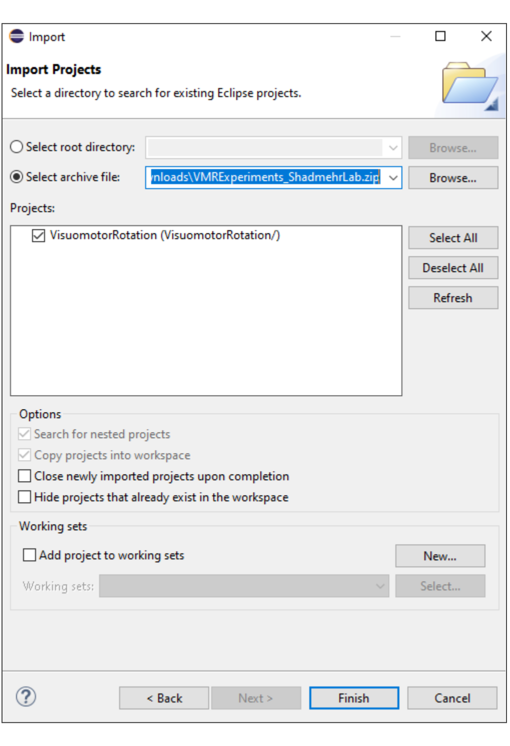


On the left side of the GUI, you should now see the Java project loaded up. This project includes both the code (under “src”), the audio file resources that were used (under “res”), and external .jar files that are needed for the program (under “bin”).

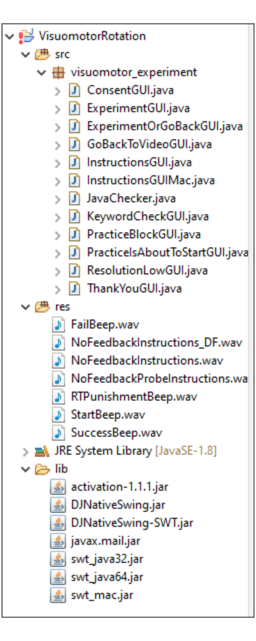


However, we are not done yet. There is a red “!”, indicating an error when we imported the code. This is due to the external .jars failing to migrate properly. To resolve this issue, right click on the parent folder titled “VisuomotorRotation,” and click “Properties.” The following window will open:

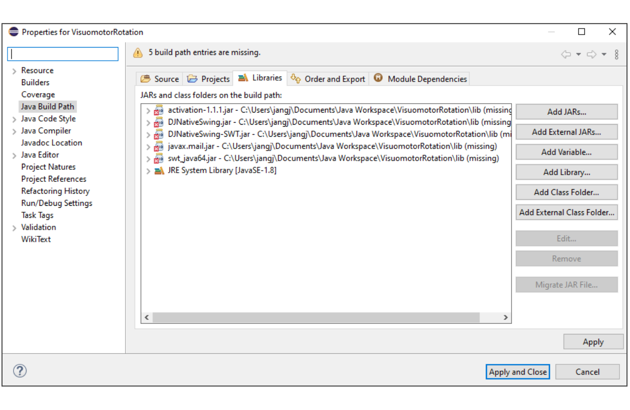

Select all of the .jar with red X’s, and click “Remove” on the right side of the window. Then, click “Add External JARs,” and navigate to the “lib” folder for this project, located within the Eclipse workspace (it is likely that the file explorer will open this folder directly).

Here, select the following files: activation-1.1.1.jar, DJNativeSwing.jar, DJNativeSwing-SWT.jar, and swt_java64.jar. You can ignore the other .jar files for now (by the time you read this manual, it is very likely that I have removed all outdated/deprecated external .jars, so you many only see swt_java32.jar as the .jar that is left over).

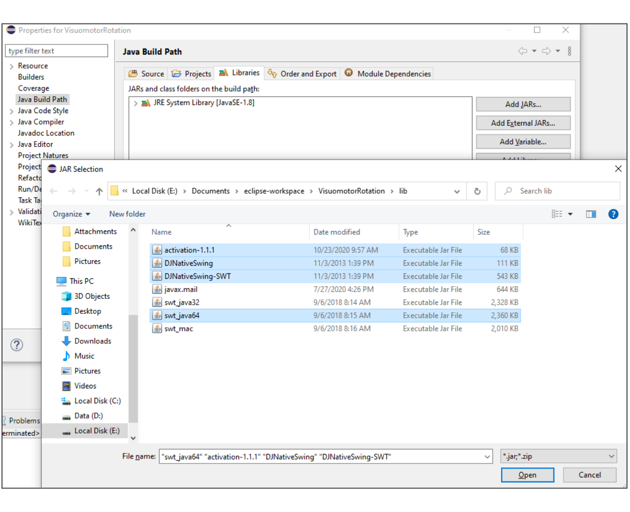

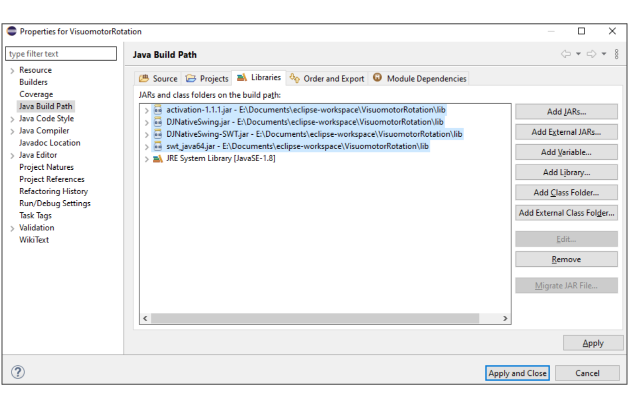

Click “Open,” then click “Apply and Close.” The errors should be resolved, and the code is now ready to run.

### Quick detour - how does the code work?

#### Overview

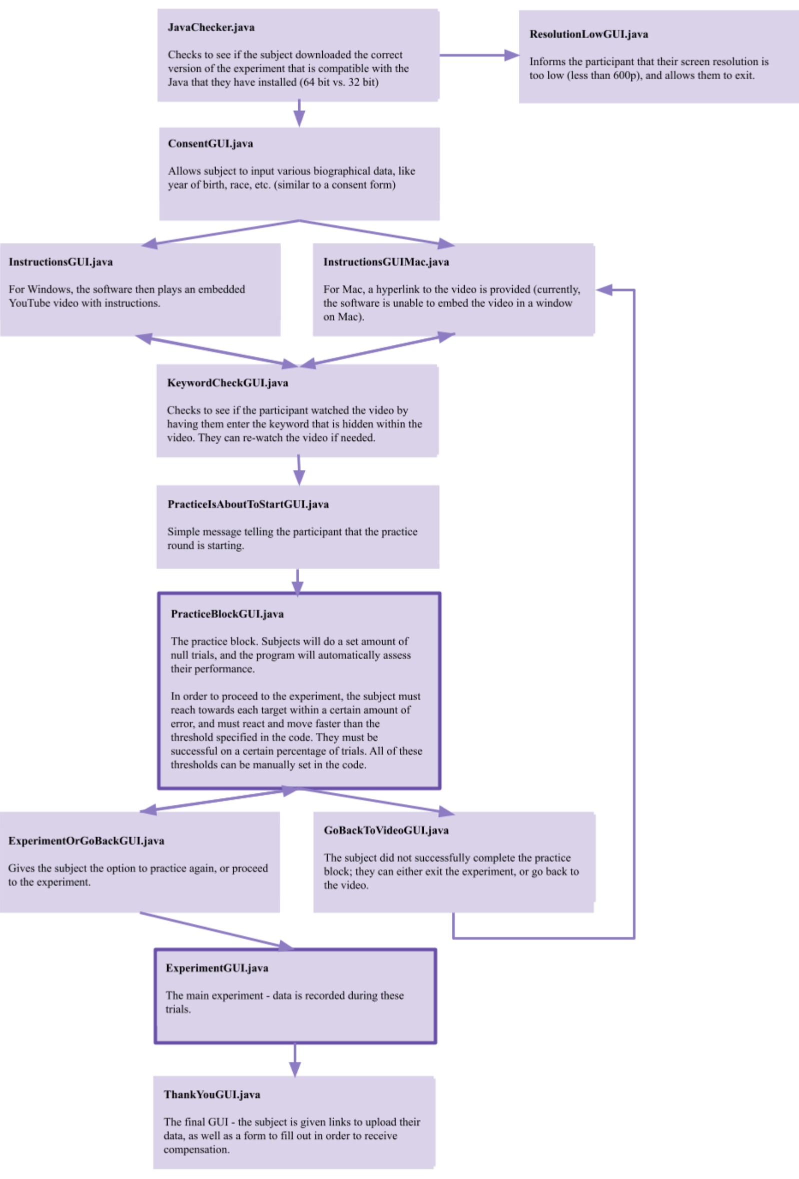


#### The experiment itself

The code begins by initializing parameters, variables, and GUI components, and designing the GUI window. Arrays containing trial data (target locations, rotation values, etc) are also calculated and generated.

Next are the MouseListener methods, which fire when the mouse enters or exit specific dots in the GUI. These methods allow the code to know when the mouse is currently within either the start or target dot.

Next are the various threads, which act similarly to a finite state machine:

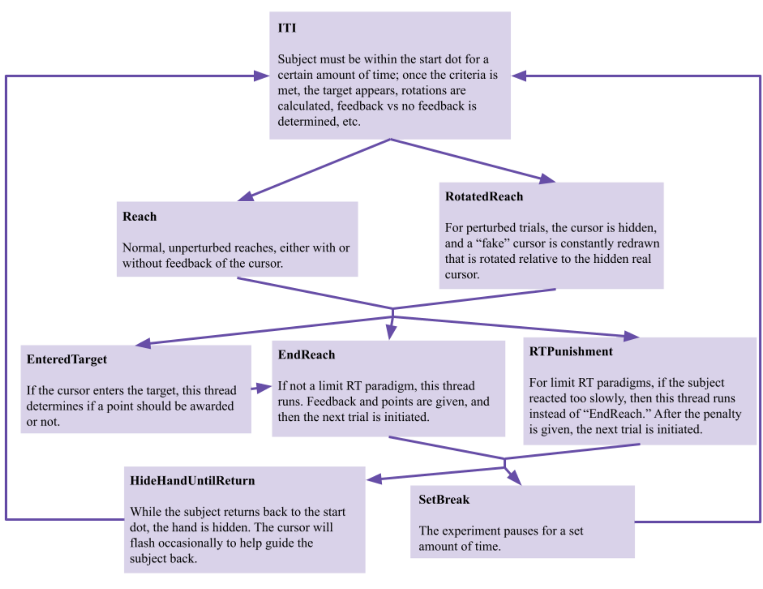


Some key points:

- Rotation perturbations are generated by hiding the real cursor, and constantly redrawing a “fake” cursor on the screen. The coordinates of this fake cursor are constantly calculated by applying a rotation matrix on the coordinates of the hidden real cursor.
- Points are scored when the cursor enters the target at the correct speed. During rotation trials, there is a second invisible and rotated target placed on the GUI with a MouseListener method attached to it. When the invisible, real cursor enters the invisible, rotated target, a point can be scored (and from the subject’s perspective, they see the fake cursor enter a visible target). A point can also be scored if the fake cursor enters the visible target on the screen, ensuring that there is still a way to score a point in case the invisible target doesn’t trigger its MouseListener method when the invisible cursor enters. Flags are added to ensure that only 1 point can be scored on a trial, no matter how many times the cursors enter their targets.

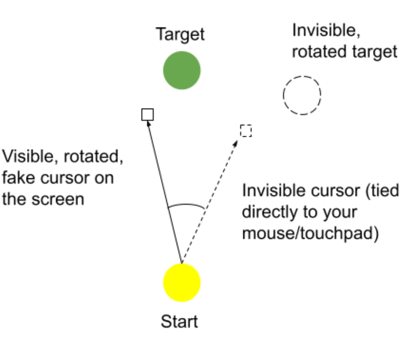


- Once the cursor crosses a specific radius around the start dot, the cursor is frozen in place and the trial ends. To calculate this freeze, the coordinates of the cursor are constantly recorded; once it crosses the radius, the coordinates of the cursor just prior to crossing the radius + the coordinates just after crossing the radius are taken, and the radius crossing point is calculated. The cursor is then displayed at this coordinate.

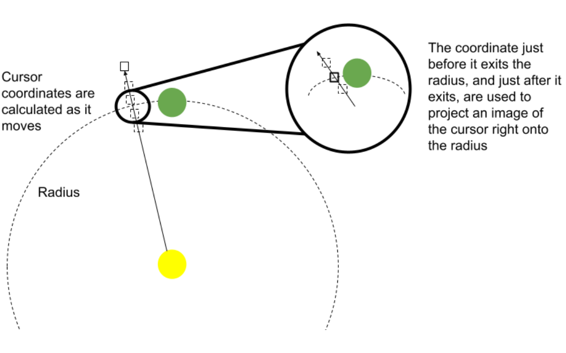


- LimitRT experiments: for these paradigms, the subject must react to the target appearance (i.e. initiate their movement) within a certain threshold; if they react too slowly, the experiment enters the RTPunishment thread, and the subject is penalized (no points awarded, a long beep sound, and a warning message). The threshold at which subjects need to react is automatically calibrated - first during the practice block, and then during the null trial period of the experiment. The calibration works as follows:
  - The first half of the practice block is designated as null trials with no limit RT condition.
  - During the second half, the code looks at a moving window of n trials; if the subject reacts successfully on y% of these n trials, then the reaction time threshold is lowered slightly. Conversely, if they reacted too slowly on many of these trials, then the threshold is increased slightly.
  - There is a lower and upper cap for this threshold, to ensure that the subject doesn’t calibrate to an unreasonably high or low threshold.
  - At the end of the practice block, the threshold value that was calibrated is saved, and then continues to be calibrated during the baseline period of the experiment itself.
  - The size of the moving window (“n”), threshold for success (“y”), threshold increment value, etc. can all be customized in the code.

○ An example:

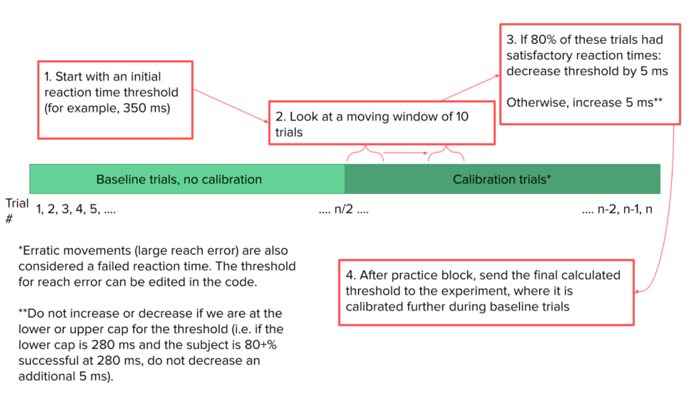


- Note: the code is commented if you are interested in learning more about a feature that is not mentioned in this manual.

### Creating an experiment

Before running the code, there are many parameters and variables that you can change to customize the content of your experiment. These are all marked by "TODO" flags so that they are easy to find. They are located in the following .java source files:

1. InstructionsGUI.java and InstructionsGUIMac.java - After creating an instructional video and uploading to YouTube, you can link this video in the Java code so that it is embedded within a GUI window.

```
48⊖     public JPanel getBrowserPanel() {
49          // TODO
50          /**
51           * You can upload an instructions video on YouTube, then place the link of the video here
52           * It will embed in a JWebBrowser and play within the experiment itself
53           * In order to ensure the video is full screen and autoplays, the URL must be in a specific format:
54           * https://www.youtube.come/embed/[unique string of characters]?autoplay=1
55           *
56           * Example: https://www.youtube.com/embed/oI0BH37DyjM?autoplay=1
57           */
58          String VideoURL = "https://www.youtube.com/embed/oI0BH37DyjM?autoplay=1"; // video for regular, gradual, and generalization experiment type
59  //      String VideoURL = "https://www.youtube.com/embed/lIuQ3nPoNTk?autoplay=1"; // video for limit RT
60  //      String VideoURL = "https://www.youtube.com/embed/frB9P4yTHKQ?autoplay=1"; // video for delayed feedback
61
```

If the URL is in the format:

https://www.youtube.com/embed/[UNIQUE VIDEO CODE]?autoplay=1

then the video will be fullscreen, and start auto playing when the subject reaches this GUI. You can find the unique video code if you watch the video in your browser and look at its URL:

youtube.com/watch?v=adLGHcj_fmA

2. KeywordCheckGUI.java - If you changed the keyword in your video, you can edit it into the code here.
3. PracticeBlockGUI.java and ExperimentGUI.java - Here, you can edit all of the parameters for both the practice block and experiment. A full list of parameters, along with explanations, are provided in the "What Do the Parameters Mean?" section of this manual.
4. ThankYouGUI.java - You can add a link to your file request here. In addition, if there is any paperwork that the subject must fill out for compensation, you can add a link to the .pdf here.

You can now run the code within Eclipse in order to see that it works. In order to run any source file of code, enter that file's code, then press the green play button at the top of the window:

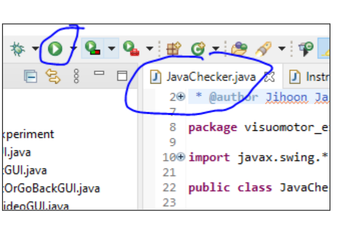

In order to stop the code, you can press the red stop button towards the bottom of the window, above the console:

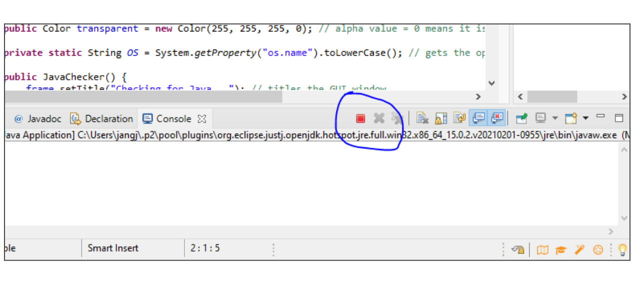

If there are any errors in your code, they will display in the console, which will allow you debug:

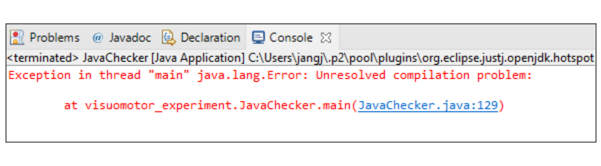


### Changing the "bitness" of your experiment

Once you are satisfied with how your experiment runs in the IDE, you can then export it into a runnable executable file. There is no need to make separate versions for Windows or Mac - the program will automatically run different blocks of code depending on the OS of the device.

However, you MAY need to create a 64 bit version (for Mac, and most Windows devices) and a 32 bit version (for a few Windows devices). Most subjects using Windows will install the latest 64 bit version of Java, but subjects using older devices or Java versions may have 32-bit and will thus require a 32 bit experiment. In order to change the bitness of the experiment, do the following:

1. In JavaChecker.java, change the String variable "bit" to either "64" or "32" (the version you are currently in the process of creating).

```
public static String bit = "64";
```

2. Then, right click on the experiment project in the menu on the left side:

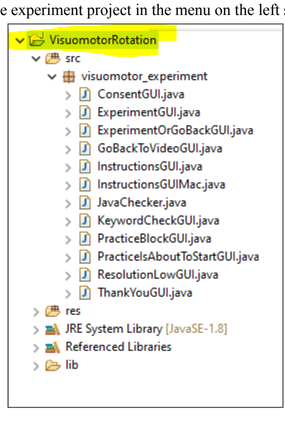

and click "Properties."

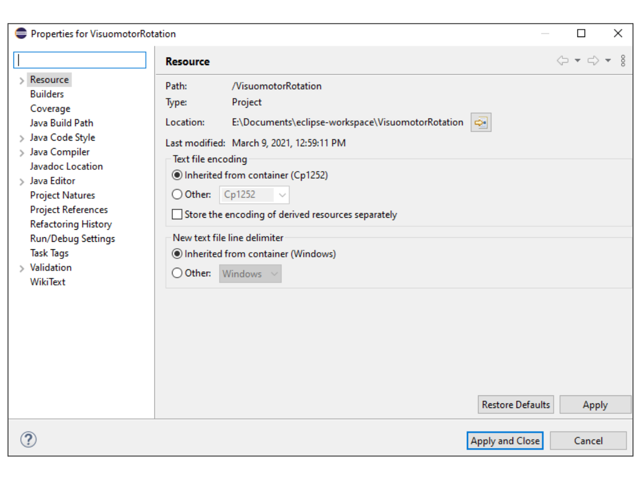

3. Navigate to "Java Build Path," and then to the "Libraries" tab.

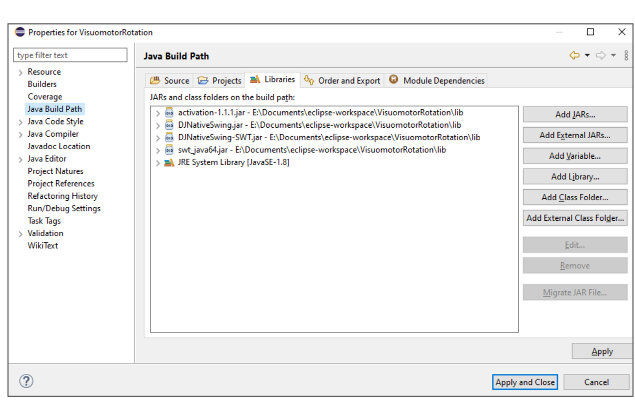

4. If you are making a 32 bit version, click "swt_java64.jar" and click "Remove" on the right side of the screen. If making a 64 bit version, do the opposite.

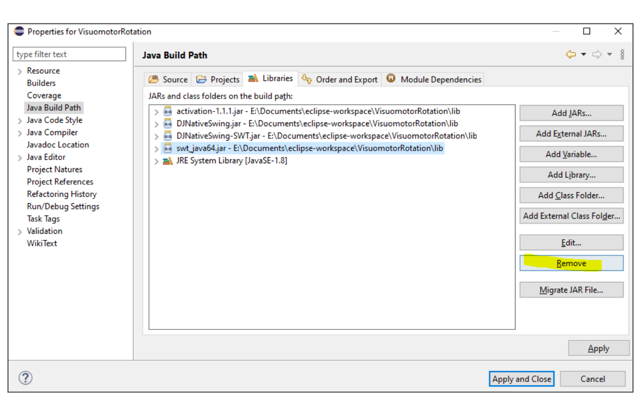

5. Then, click "Add External JARs," navigate to the "lib" folder, and select the correct swt_java file (if making a 32 bit program, choose swt_java32.jar, and vice versa).

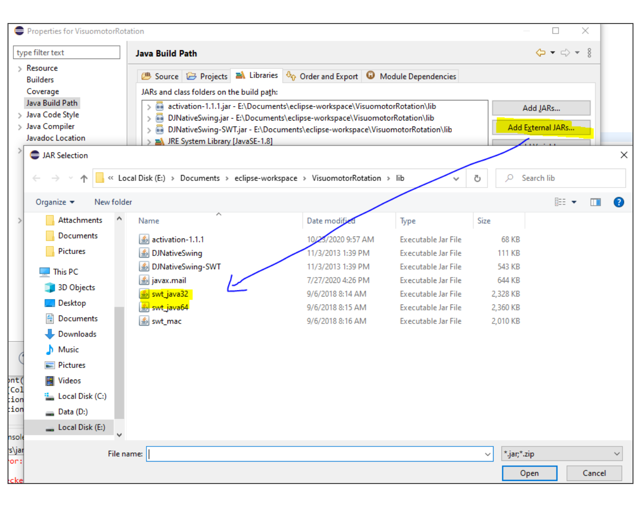

6. You can then press "Apply and Close," and make sure one last time that the bitness of the swt_java file AND the String variable from step 1 both match.

### Exporting the experiment

In order to export the program as an executable file, right click on the project in the menu on the left side, and click "Export."

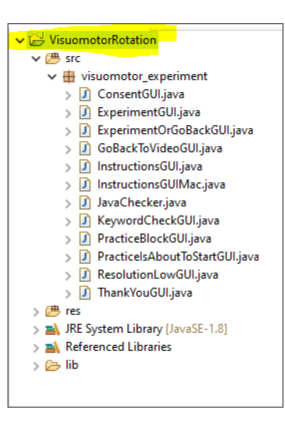

Select the "Java" folder, choose "Runnable JAR file," and click "Next."

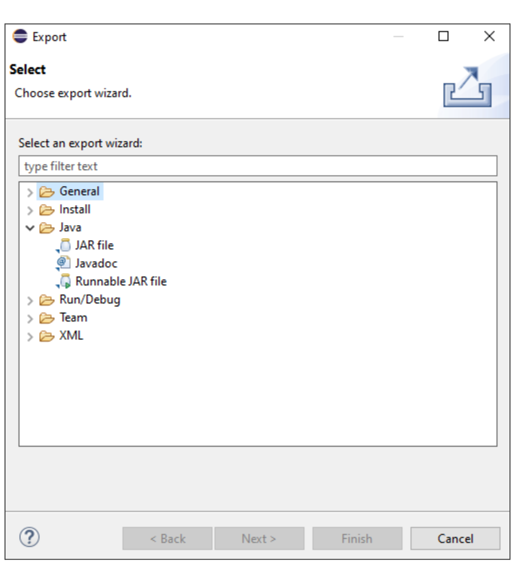

Under "Launch configuration," choose "JavaChecker." This will ensure that the entry point of the executable (the first block of code that is run) when the subject double clicks on the file is the JavaChecker code.

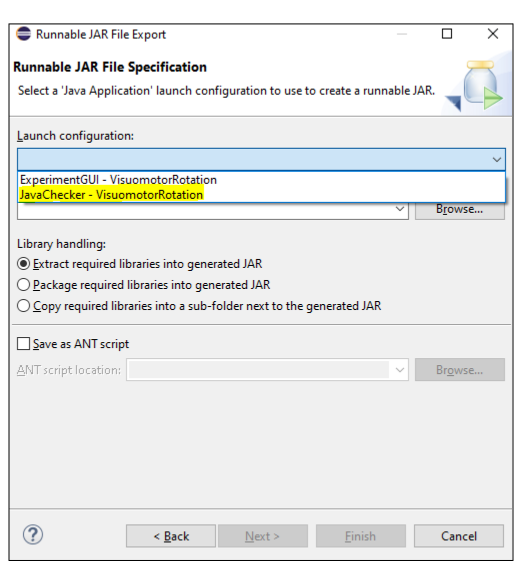

You can then click "Browse" to choose an Export destination in your file system, as well as name your executable .jar file.

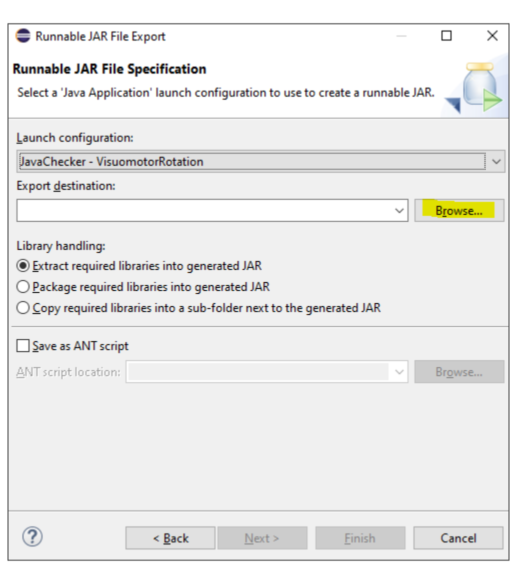

Press "Finish," and the executable .jar file will generate.

If you are testing something out and want to view error messages while running the executable file, you can open it via the command line (not double clicking). Error messages will print in the command line. Navigate to the folder with the executable file and enter the following command:

```
java -jar [NAME OF EXECUTABLE].jar
```

### Analyzing data

Every time the experiment runs, it will output a unique text file containing the data. Using the programming language of your choice, you can analyze this data by loading in the text file and parsing it. For example, in Matlab this can be achieved using the "regexp" function to look for specific regular expressions.

These last two parameters are only found in the practice block:

- **angle_threshold** - Defines the threshold of reach error for a movement to determine if it is erratic or not. This field should be a double.
- **practice_percent_correct_threshold** - Defines the percent of trials during practice that must be acceptable (reacted quickly, reached quickly, not erratic) to proceed. This field should be a double.
